# *SHOOTMERISTEMLESS* autoactivation: A prerequisite for fruit metamorphosis

**DOI:** 10.1101/2024.02.23.581830

**Authors:** Yang Dong, Zhi-Cheng Hu, Mateusz Majda, Hao-Ran Sun, Yao Zhang, Yi-Ning Ding, Quan Yuan, Tong-Bing Su, Tian-Feng Lü, Feng Gao, Gui-Xia Xu, Richard S. Smith, Lars Østergaard

## Abstract

In animals and plants, organ shape is primarily determined during primordium development by carefully coordinated growth and cell division^1-3^. Rare examples of post-primordial change in morphology (reshaping) exist that offer tractable systems to study mechanisms required for organ-shape determination and diversification. One such example is the heart-shape formation of *Capsella* fruits that occurs by reshaping the ovate spheroid gynoecium upon fertilization^4^. Here we use whole-organ live-imaging to show that dynamic changes in growth and cell division coupled with local maintenance of meristematic identity drives *Capsella* fruit-shape formation. At the molecular level, we reveal an auxin-induced mechanism ultimately descending on a single *cis* regulatory element to mediate morphological alteration. This element resides in the promoter of the *Capsella rubella SHOOTMERISTEMLESS^5^* (*CrSTM*) gene. The CrSTM meristem identity factor positively regulates its own expression through binding to this element thereby providing a feed-forward loop at the position and time when protrusions emerge to form the heart. Independent evolution of the STM-binding element in *STM* promoters across Brassicaceae species correlates with those undergoing a gynoecium-to-fruit metamorphosis. Accordingly, genetic and phenotypic studies showed that the STM-binding element is required to facilitate the shape transition and reveals a conserved molecular mechanism for organ morphogenesis.

## Main text

Multicellular organisms produce organs with specific functions for resource acquisition and reproduction^6^. Evolution of such specialized functions is intricately linked with organ morphology diversification that are adaptive in distinct ecological niches^7-9^. For example, evolutionary radiation of fruit shape has led to reproductive strategies adapted to optimize success of the next generation^4,10,11^. Such phenotypic diversity requires modulation in the coordination of cell division and growth in reproductive structures^1-3^. However, the molecular mechanism underlying this is poorly understood.

### Early growth dynamics determine fruit shape

Members of the *Capsella* genus produce fruits that are shaped like a heart^4^. Upon fertilization of the *Capsella* flower, the female reproductive organ undergoes metamorphosis from an oblate spheroid (disc-shaped) gynoecium to a flat, heart-shaped fruit^4,12^. This shape is adopted as the two valves (seed pod walls) each form one half of the heart protruding laterally from the central replum. The *Capsella* fruit therefore provides an excellent system to study the cellular and molecular basis of organ shape establishment^12^. Using the plasma membrane marker line, *pUBQ10:acyl-YFP*^13^, we performed a large-scale analysis including a total of 18,656 cells of the female reproductive organ from developmental stage 12 (fertilization) to stage 14, during which the heart-shape of the fruit gradually emerge^4,12,14^.

Using lineage tracking maps, we quantified cell growth parameters that affect form, including growth (rate of cell area expansion), anisotropy (direction of cell expansion), and cell proliferation (Fig. 1). Analysis of cell size revealed a gradient with larger cells predominantly located at the base of the fruit and progressively smaller cells observed towards the tip (Fig. 1a,b). Interestingly, this cell-size gradient was less obvious at the stage of fertilization (Fig. 1a) and may suggest an apical-basal gradient in cell differentiation. While cell division was found evenly throughout the fruit at stage 13 (Fig. 1c), this indicates that cells at the apical part retain the competency to divide longer than the basal region. It is worth noting that this analysis also shows that – in contrast to post-fertilization growth of the *Arabidopsis* fruit^15^ – cells of the *Capsella* fruit continue to divide after fertilization (Fig. 1c). Consistently, a region with high level of growth anisotropy (directed cell growth) was observed at the apical part of the fruit growing towards a region of smaller cells at the tip (Fig. 1d). In summary, this organ-wide cellular analysis suggests that coordination of locally controlled anisotropic growth and changes in cellular growth and division are required for the reshaping of the *Capsella* gynoecium at the onset of fruit development.

**Fig. 1.**
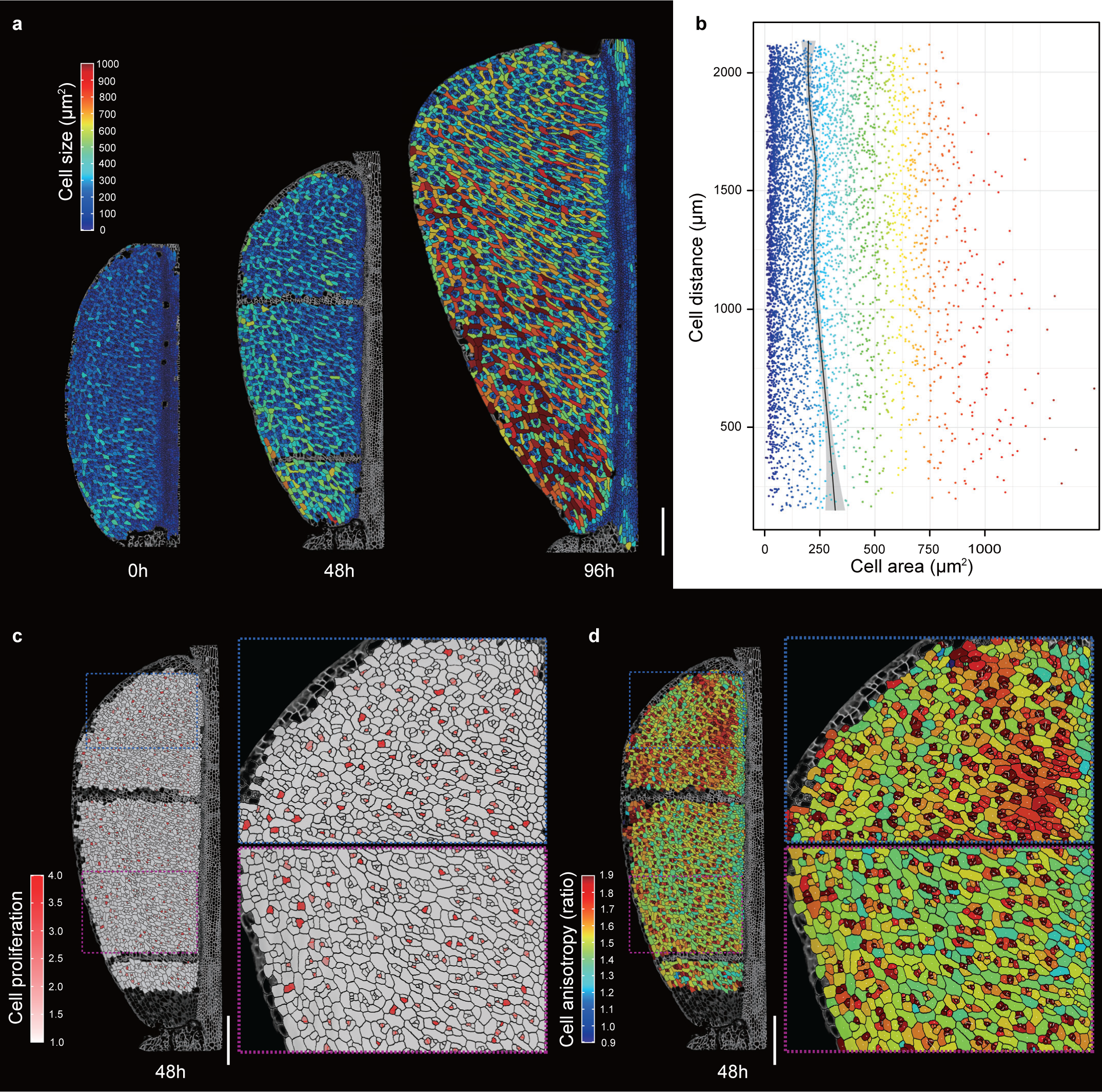
Dissecting the cellular basis of fruit-shape determination in *Capsella*. **a**, Heat-maps depicting cell sizes in the *Capsella* fruit in a 96-hour time-lapse live imaging analysis corresponding to developmental stages 12 to 14. Cell size is quantified by cell area (μm²). **b**, Scatter plot representing cell size distribution along the vertical axis of the fruit at the 96-hour time point. The Y-axis denotes inter-cellular distance, where ∼500 μm corresponds to the fruit bottom and ∼2000 μm represents the fruit top. The X-axis indicates cell size. Each dot represents an individual cell, color-coded according to the size as per the heat-map. The average cell size and confidence interval are depicted by a black line and grey shade, respectively. **c**, Heat-map demonstrating cell proliferation at the 48-hour time point in the *Capsella* fruit. **d**, Heat-map indicating cell anisotropy at the 48-hour time point in the *Capsella* fruit. Scale bars in **a**, **c**, **d**, 200 μm.

### Meristematic cell identity is maintained during metamorphosis

To characterize the developmental trajectories of the valve tips at the single-cell level, we conducted a comparative single-cell RNA sequencing (scRNA-Seq) analysis of the valve tips of stage-13 (shape change initiates) and stage-14 fruits (shape change is achieved). A total of 32,126 cells from stage-13 valve tips (∼260 fruits) and 38,349 cells from stage-14 valves tips (∼200 fruits) were isolated and cell walls removed (Extended Data Fig. 1). These cells were then subjected to scRNA-seq using droplet-based technology on the commercial 10 X chromium platform (10 X genomics) (Extended Data Fig. 1 and 2). After aligning the data to the *Capsella* genome^16^, removal of low-quality cells and filtering out genes induced during protoplast isolation (see Methods), we obtained gene expression matrices containing 17,979 genes across 17,453 cells for stage-13 samples and 18,449 genes across 23,176 cells for stage-14 samples, respectively (see Methods and Extended Data Fig. 3a, b; Extended Data Sheet 1). The analyses of the two replicates revealed a high level of consistency between replicates (Extended Data Fig. 4). We subsequently used principal component analysis (PCA) on a gene expression matrix across 2000 variable genes and identified 50 statistically significant principal components (*p*<0.05). These Principal Components were then processed with Seurat, resolving 16 and 15 distinct clusters for stage 13 and 14, respectively (Fig. 2a,b and Extended Data Fig. 5), revealing heterogeneous cell populations. The stage-13 clusters contained 158 to 3136 cells and stage-14 clusters contained 301 to 4287 cells (Extended Data Fig. 3c, d). These clusters contained similar number of cells from each replicate and the gene expression profiles exhibited a high correlation across the samples and bulk RNA-seq (Stage 13, ρ= 0.9273; Stage 14, ρ= 0.9206), demonstrating the reproducibility of the sample (Extended Data Fig. 6).

**Fig. 2.**
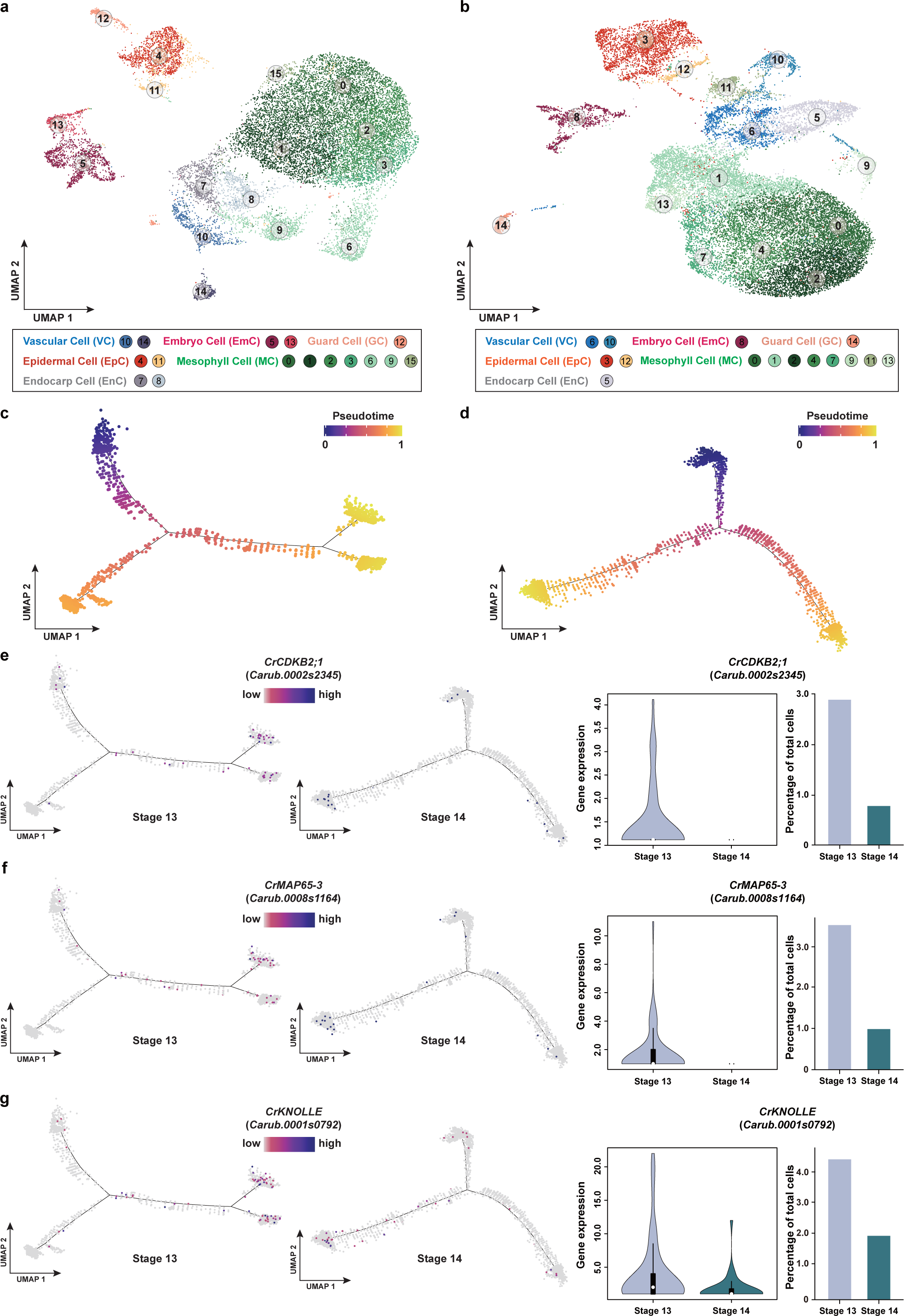
Comparative analysis of cell differentiation trajectories in the fruit valve tip epidermal cells. **a**, UMAP visualization of 16 cell clusters in stage-13 fruit valve tips (*n* = 17,453 cells). **b**, UMAP visualization of 15 cell clusters in stage-14 fruit valve tips (*n* = 23,176 cells). In **a** and **b**, each dot denotes a single cell and colors denote corresponding cell clusters, respectively. All the cell clusters can be grouped into six cell types (see also Extended Data Fig. 8) with corresponding colors, vascular cell (VC), embryo cell (EmC), guard cell (GC), epidermal cell (EpC), mesophyll cell (MC), endocarp (EnC). **c** and **d**, Monocle 2 analysis showing differentiation trajectory colored by pseudotime of the valve tip epidermal cells from stage-13 fruit (**c**, *n* = 1,775 cells, see Methods) and stage-14 fruit (**d**, *n* = 3,000 cells, see Methods). **e**-**g**, Comparisons of the expression pattern, expression level and the percentage of cells expressing the genes involved in cell cycle and cytokinesis (*CrCDKB2;1* (**e**), *CrMAP65-3* (**f**), and *CrKNOLLE* (**g**)) in the valve tip epidermal cells between stage-13 and stage-14 fruits. The gene expression level is calculated by the ggboxplot function (R ggpubr package). Each dot in the differentiation trajectory indicates a single cell.

The clusters were annotated using orthologues of *Arabidopsis* genes with known biological functions and expression patterns in addition to known markers identified in other scRNA-Seq datasets^17^ (Extended Data Sheet 1 and 2; Extended Data Fig. 7). Based on this analysis, we could group the clusters into six individual cell types at both developmental stages (Fig. 2a,b), which is consistent with the anatomical structures of the fruit valves (Extended Data Fig. 8). The live imaging results 1 suggest a progression of cell differentiation in valve epidermal cells (EpCs) at the tips. We therefore compared the developmental trajectories of the valve tip EpCs during the shape transition between stage 13 and 14 (Fig. 2c,d). The EpC cluster was identified as they specifically express marker genes, such as ATML1, FDH and DCR^18-20^ (Extended Data Fig. 9 and Extended Data Sheet 2). Pseudotime analysis by ordering cells along a reconstructed differentiation trajectory using Monocle2 shows that genes associated with cell cycle (*CYCLINS*, *CDKs* and *cytokinesis*) are expressed more in stage-13 than in the stage-14 samples (Fig. 2e-g; Extended Data Fig. 10; Extended Data Sheet 2). Interestingly, comparison of the developmental trajectories of the EpCs revealed that both the expression level of cell-cycle genes and the proportion of expressing cells were significantly reduced from stage 13 to 14, suggesting a progressive decrease in cell division activity (Fig. 2 e-hand Extended Data Fig. 10).

To validate the scRNA-Seq results, we generated GUS-reporter lines for two cyclin genes, *CrCYCB1;1* and *CrCYCB1;2* and found that they are expressed in valve tips during shape formation and reduced in this region later in development (Extended Data Fig. 11). Moreover, application of flavopiridol (FVP), a cyclin-dependent kinase (CDK) inhibitor that blocks the cell cycle progression at the G1-S and G2-M phases^21^, significantly suppress formation of the heart-shaped fruit by reducing the growth of the valve tips (Extended Data Fig. 12). Consistent with the observation that more cells undergo cell division in the valve tips at stage 13 compared to stage 14, fruits at stage 13 were more affected by FVP than those at stage 14 (Extended Data Fig. 12).

Taken together, the scRNA-Seq analysis provides transcriptomic support to the cellular growth data from the live-imaging analysis that cells in the valve tip maintains meristematic identity during the transformation in shape from gynoecium to the fertilized fruit.

### *CrSTM* autoregulation controls *Capsella* fruit shape

Previously, we showed that the valve tips have high auxin levels resulting from localized auxin biosynthesis^12^ coinciding with meristematic activity as illustrated here by live imaging and correlating with the scRNA-Seq analysis. This pattern of cell division and auxin dynamics is analogous to the Shoot Apical Meristem (SAM), where new organ primordia are initiated in a regular pattern through the formation of local auxin maxima and active cell division^22,23^. In *Arabidopsis*, stem cell identity is maintained by an intricate network of key regulators including transcription factors such as *WUSCHEL* (*WUS*), *SHOOTMERISTEMLESS* (*STM*), *KNAT1* (also known as *BREVIPEDICELLUS*), *KNAT2* and *KNAT6*^5,24-26^. We speculated that specific expression of these meristem-identity genes in the valve tips of could be responsible for delaying differentiation in this region. To this end, we created GUS reporter lines for *Capsella* orthologues of these genes. Whereas they were all expressed in the SAM, only *CrSTM* revealed expression during fruit development (Extended Data Fig. 13). Interestingly, a dynamic expression pattern of *CrSTM* coincided with the progressive establishment of the heart-shaped fruit from stage 12 to stage 14 (Fig. 3a-d and Extended Data Fig. 13). In stage-14 fruits, *pCrSTM:GUS* expression is specifically localized in the valve tips (Fig. 3d), overlapping with the expression of auxin biosynthesis genes (*pCrYUC9:GUS*, *pCrTAA1:GUS*), and the *pDR5:GUS* auxin signaling reporter^12^. These data therefore suggest a potential role of *CrSTM* in regulating shape formation during *Capsella* fruit development.

**Fig. 3.**
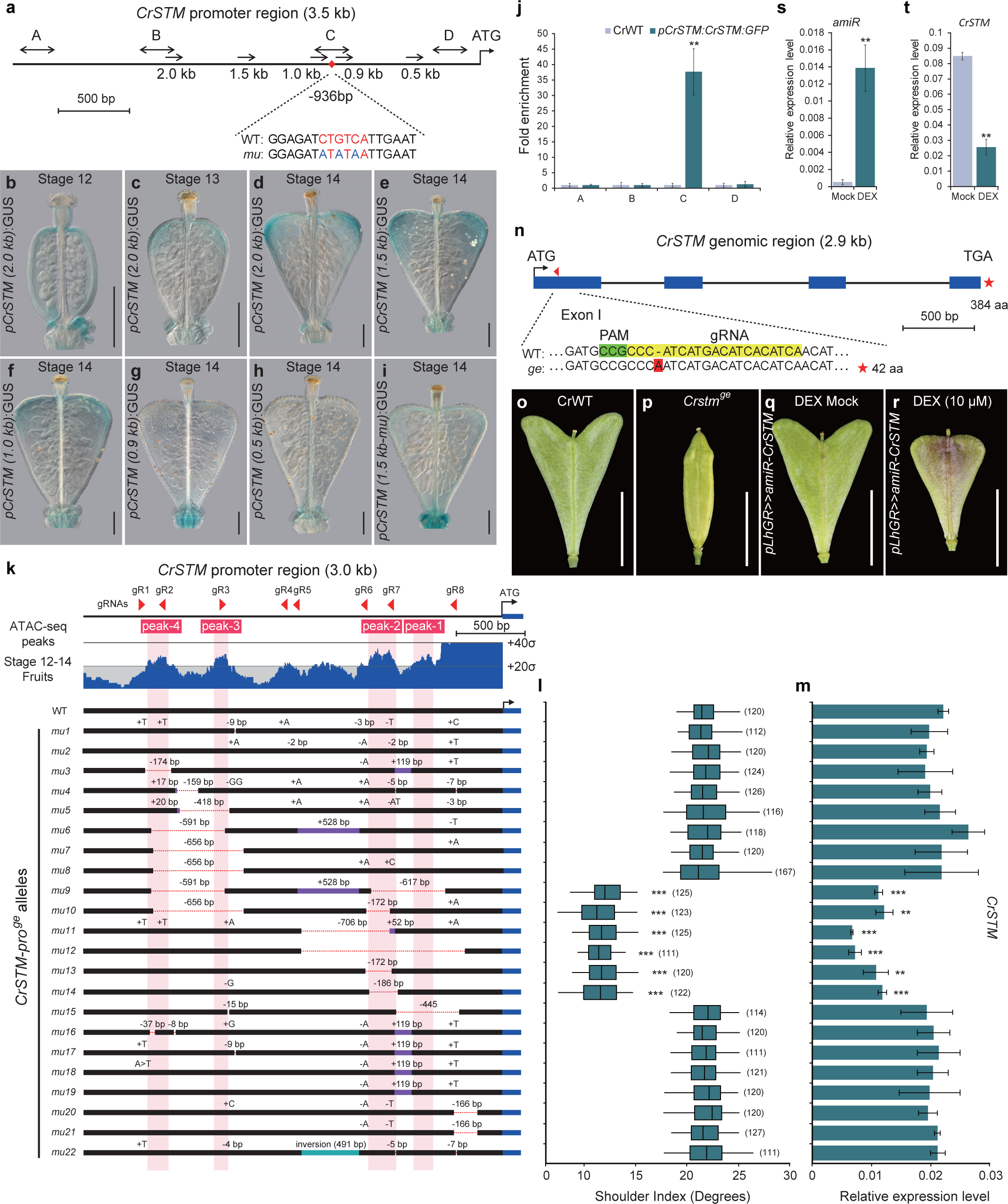
Functional analysis of the *CrSTM* locus reveals a positive feed-forward regulatory loop. **a**, Schematic of the 3.5-kb *CrSTM* promoter upstream of the start codon. The arrows indicate the promoter deletion series in the GUS reporter analysis in **b** to **i**. The double arrows indicate the regions used for ChIP analysis. The red diamond indicates the position of the STM-binding site (-936 bp, CTGTCA red letters), and the mutated version (mu, ATATAT) used in the *1.5 kb* GUS reporter construct is aligned below. **b**-**d**, *CrSTM* expression pattern in the *Capsella* fruits shown by GUS staining at stage 12 (**b**), stage 13 (**c**) and stage 14(**d**). **e**-**i**, *CrSTM* promoter activity analysis shown by GUS staining using 1.5-kb (**e**), 1.0-kb (**f**), 0.9-kb (**g**), 0.5-kb (**h**) and 1.5-kb-mutated (**i**) *promoter:GUS* reporter lines. **j**, ChIP analysis of CrSTM:GFP protein associated with the *CrSTM* promoter (region C in **a**). **k**, ATAC-Seq coverage (blue) on the 3.0-kb *CrSTM* promoter region (upper panel) using stage 12-14 fruit samples. The grey lines mark the mean and standard deviation. ATAC-seq peaks are numbered sequentially from ATG and shaded in light pink. The red triangles indicate the gRNAs used for generating promoter-editing alleles. The lower panel shows the 22 *CrSTM* promoter alleles. The deletions are indicated by dashed red lines, and the insertions and inversions are shown by purple and green rectangles, respectively. **l**, Quantification of fruit shape by shoulder index (see also Extended Data Fig. 12). The number of fruits used for quantification is indicated. For each allele, at least three plants were sampled. **m**, Analysis of *CrSTM* expression from stage 13/14 fruits of each allele. Error bars represent the SD of three independent biological replicates. **n**, Schematic of the *CrSTM* genomic region (2.9 kb) with exons and introns indicated by blue rectangles and black lines, respectively. The single gRNA targeting the first exon is indicated by a red triangle. The genotyping result is presented below the graph. **o** and **p**, Fruit morphology of CrWT (**o**) and *Crstm^ge^*(**p**) at stage 17. **q** and **r**, Fruit morphology of Mock (**q**) and DEX (**r**) treatment of the *pLhGR>>amiR-CrSTM* lines at stage 17. **s** and **t**, Expression analysis of the amiR (**s**) and endogenous *CrSTM* transcripts (**t**) in stage 13/14 fruit samples upon DEX treatment. Scale bars, **b**-**i**, 100 μm; **o**-**r**, 5 mm. **, p < 0.01; ***, p < 0.001 (Student’s t test).

*CrSTM* encodes a class-I KNOX transcription factor that controls plant growth and development by maintaining the stem cell population in the SAM^5,27^. Regulatory variations in the *STM* gene and its homologs is required to control axillary meristems and complex leaf shapes^28,29^. *STM* has also been implicated in the regulation of leaf-shape diversity affecting patterns of growth proliferation and differentiation^30^. We next made an attempt to dissect the functional regions that instruct *CrSTM* to function in fruit-shape determination. We generated a series of promoter deletions and narrowed the functional region required for valve tip expression of *CrSTM* down to a 93 bp segment around 1000bp upstream from the transcription start site (Fig. 3a,e-h and Extended Data Fig. 14). Within this region, we identified a hexamer, CTGTCA, previously reported as the recognition sequence for STM in *Arabidopsis*^31^. Intriguingly, this sequence is not present in the *AtSTM* promoter in the orthologous region (Extended Data Fig. 15) and mutating it in the *pCrSTM(1.5k-mu):GUS* reporter line abolished expression in the valve tips (Fig. 3a,i). By Chromatin Immuno-Precipitation (ChIP), we found that a CrSTM-GFP fusion associated with a fragment containing the STM element (Fig. 3j). In addition, using a dexamethasone (DEX)-inducible version of *CrSTM,* we could induce *pCrSTM(1.5k):GUS* expression and show that this depended on the presence of the CTGTCA element (Extended Data Fig. 16a,b). Moreover, DEX-induced activation of *CrSTM* expression was unaffected by the presence of the protein synthesis inhibitor, cycloheximide (CHX) (Extended Data Fig. 16c). These experiments demonstrate that *CrSTM* is autoregulated and sustains expression in the fruits through a positive feedback regulatory loop. This finding is furthermore supported by an Assay for Transposase-Accessible Chromatin using Sequencing (ATAC-Seq) experiment where increased chromatin accessibility in the region around the CTGTCA element was found to be associated with temporal *CrSTM* expression in the fruit (Fig. 3k and Extended Data Fig. 17).

To further assess the functional importance of the STM binding site in fruit-shape determination compared to other regions of the *CrSTM* promoter, we created 22 gene-edited lines with diverse deletions in their *CrSTM* promoter region using a CRISPR/Cas9-based gene editing system with eight guide RNAs (gRNAs) distributed across a 3-kb region upstream of the *CrSTM* coding region (Fig. 3k and Extended Data Fig. 18). We evaluated the effects of the variation by quantifying the fruit shape with shoulder index^12^ and *CrSTM* expression level (Fig. 3l,m). We found that deletions involving gRNA6 and gRNA7 (*mu9-mu14*) generated plants with compromised *CrSTM* expression and defects in valve tip development (Fig. 3k-m and Extended Data Fig. 19). These deletions share a ∼150-bp region, which includes the STM binding site, further supporting the pivotal role of this element in fruit-shape determination with little contribution from other regions of the *CrSTM* promoter.

The autoregulatory expression of *CrSTM* raises the question as to how it is initially kicked off. It was previously shown that *STM* in *Arabidopsis* is regulated partly by Auxin Response Factors (ARFs) in the floral meristem^32^. Given that auxin dynamics at the valve tips plays a critical role in heart-shape formation^12,14^, it is possible that *CrSTM* expression is affected by auxin. Two ARFs are particularly important for fruit growth in *Arabidopsis*, namely ARF6 and ARF8^33^ and are therefore candidates for regulating *CrSTM* expression. We identified the closest orthologs of *ARF6* and *ARF8* in *Capsella* (*CrARF6* and *CrARF8*) and created single and double knock-out mutants by gene editing to assess their role in *Capsella* fruit development. While neither *Crarf6^ge^* nor *Crarf8^ge^* single mutant exhibit a defect in fruit valve development, the *Crarf6/8^ge^* double mutant produced valves that failed to grow after fertilization (Extended Data Fig. 20). Interestingly, *CrSTM* expression was significantly reduced in the *Crarf6/8^ge^* double mutant but not in the either single mutant (Extended Data Fig. 21), and ChIP experiments with GFP fusions showed that both CrARF6 and CrARF8 associate with regions of the *CrSTM* promoter (Extended Data Fig. 21). These data suggest auxin initiates *CrSTM* expression in the fruit through the redundant function of *CrARF6/8*. Once activated, *CrSTM* sustains its expression via autoregulation, thus delaying cell differentiation and regulating the morphological change.

To directly test the requirement for *STM* in fruit-shape formation, we generated knock-out lines of *CrSTM* using a gene editing approach targeting Exon 1 of the *CrSTM* gene (Fig. 3n). While many of these lines expectedly showed severe phenotypic defects from an early stage as described for *stm* mutants in *Arabidopsis*^5^, some lines produced inflorescences and fruits (Extended Data Fig. 22). Strikingly, these fruits completely lacked outgrowth of the valve tips and failed to undergo the characteristic change in shape from the disc-shaped gynoecium to the heart shape of fertilized fruits (Fig. 3o, p and Extended Data Fig. 23). These *Crstm^ge^* mutant fruits failed to set viable seeds, precluding us from further studying its contribution to fruit development. To overcome this, we generated a transgenic line expressing a DEX-inducible artificial microRNAs (amiR) that targets the first exon of *CrSTM* (*pLhGR>>amiR-CrSTM*). Upon DEX treatment, microRNA expression was induced and endogenous *CrSTM* mRNA levels were down-regulated (Fig. 3s,t). Down regulation of *CrSTM* led to production of smaller fruits lacking valve tips, thus abolishing the heart shape (Fig. 3q,r and Extended Data Fig. 23). The phenotype of DEX-treated *pLhGR>>amiR-STM* transgenic fruits is therefore very similar to the FPV-treated fruits (Extended Data Fig. 12), suggesting that down-regulation of *CrSTM* decrease cell division in the valve tips, which in turn perturbs the fruit-shape development program in *Capsella*. In agreement with this, expression of genes involved in cell cycle control was significantly reduced upon DEX treatment (Extended Data Fig. 24). This experiment demonstrates that the *Capsella STM* orthologue is required for the change in fruit shape from gynoecium to mature fruit.

Finally, we created a version of the *Arabidopsis STM* gene engineered to include the STM binding site at the equivalent position in the promoter as in the *CrSTM* gene (Extended Data Fig. 25a). A ChIP experiment showed that an AtSTM-GFP tag indeed associate with this engineered region (Extended Data Fig. 25b). While gynoecia from transgenic *Arabidopsis* lines expressing this construct showed no morphological difference compared to wild type, their fruits exhibited a shape change and over-proliferation of the epidermal cells (Extended Data Fig. 25c). These data demonstrate that acquisition of the STM-binding element in the *AtSTM* promoter induces ectopic *STM* expression and cell proliferation leading to a morphogenetic change, showing that STM autoregulation is sufficient for fruit metamorphosis.

### A conserved process for morphological diversity

The Brassicaceae family contains more than 3800 species and is characterized by a diversity of fruit shapes that are derived from an ancestral cylindrical shape^34,35^. To test if the STM binding site exists in other *STM* orthologous genes, we analyzed the promoter sequences of a panel of 43 Brassicaceae species with *Cleome violacea* (Capparaceae) as the outgroup (Fig. 4a). We found that in the equivalent region of the *STM* promoter, the STM-binding site has evolved repeatedly within the Brassicaceae family (Fig. 4a). Remarkably, we noticed a perfect correlation between species harboring the STM-binding site and those undergoing a metamorphosis from the gynoecia to the mature fruit stage (Fig. 4b). We hypothesize this could be due to post-fertilization cell divisions in the fruits caused – at least in part – by *STM* expression and autoregulation. In support of this proposal, our molecular evolution analysis, moreover, revealed that the STM-binding site in Brassicaceae has been under strong selection (Extended Fig. 26), further substantiating the pivotal evolutionary role of this particular 6-bp sequence in adaptation.

**Fig. 4.**
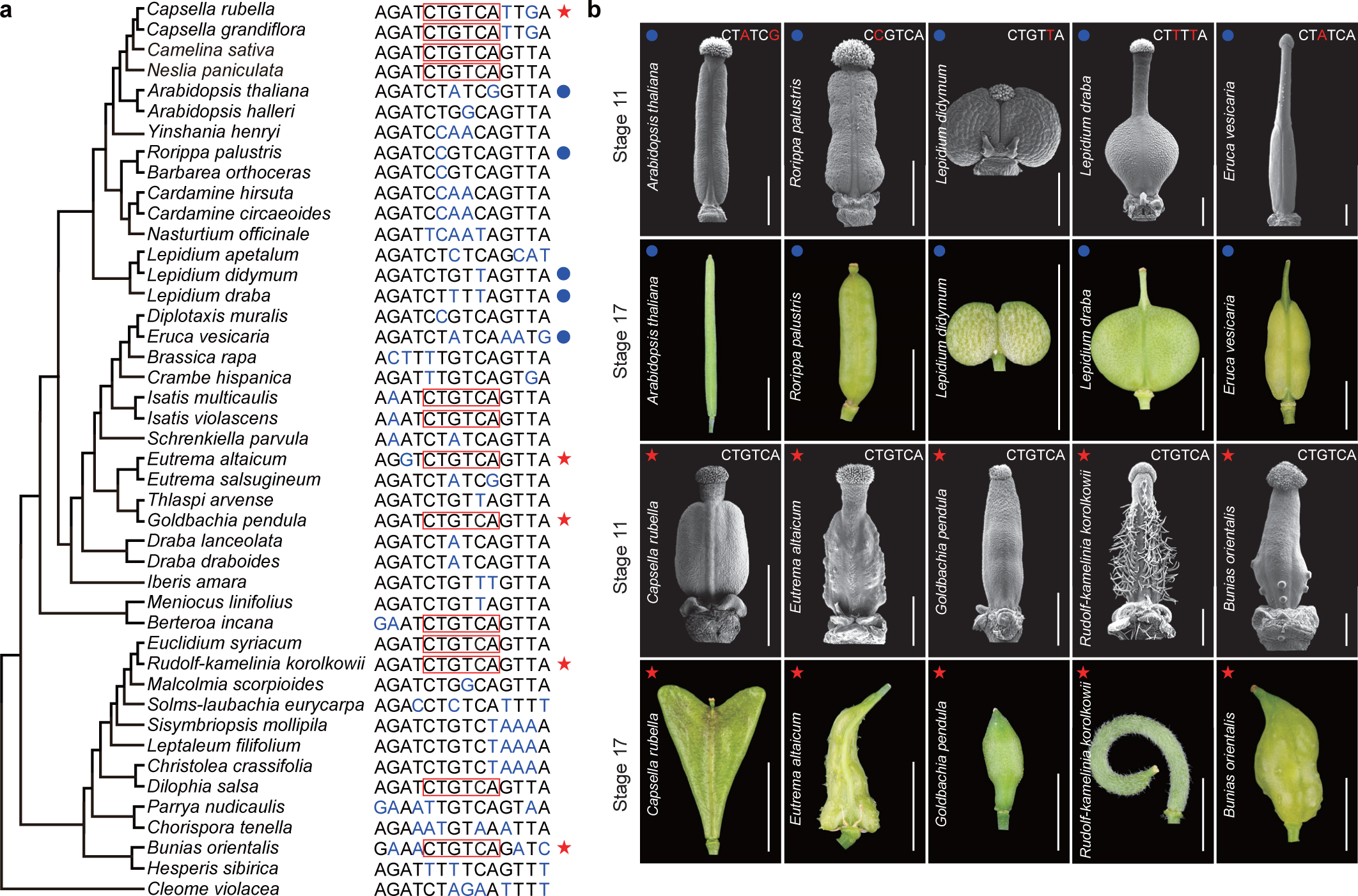
Evolution of the STM-binding sites associated with fruit-shape diversification in Brassicaceae. **a**, Phylogenetic tree of 43 Brassicaceae species with *Cleome violacea* (Capparaceae) as the outgroup. The relationship between species was reconstructed based on published phylogenetic studies^35-37^. The orthologous region of the STM-binding was aligned. The red boxes indicate the consensus STM-binding site. The conserved sequences are shown in black and the diverged sequences are shown in blue, respectively. The representative species with (red stars) or without (blue dots) the STM-binding site selected for morphological analysis are indicated on the right side. **b**, Scanning electron microscopic (SEM) graphs of the gynoecia at stage 11 and whole-mount fruit images at stage 17 of Brassicaceae species with (red stars) or without (blue dots) STM-binding site shown in **a**. The WT and mu version of STM-binding sites of each species were shown in the SEM images. Scale bars, 400 μm (SEM images); 5 mm (whole-mount images).

Nature is rich in morphological diversity, not least among fruits from flowering plants exhibiting an immense variation in shape. Our work shows that fruit shape depends on specific local patterns of cell division and cell growth. This process involves a positive and autoregulatory loop of the meristem-identity factor, STM, centered around the presence or absence of an STM binding site in the promoter of the *STM* gene itself. The mechanism is initiated by local auxin-induced signaling and is required for the metamorphic shape change occurring between gynoecium and fruit formation. While great diversity in fruit shape exists across species independently of this mechanism (Fig. 4b), STM autoregulation relieves the morphological constraint imposed by the gynoecium shape. Since autoregulation of STM appears to have been co-opted independently multiple times within the Brassicaceae family, it is possible that selection pressure for obtaining the STM binding site is strong in the evolutionary history of morphological diversity.

## Methods

### Plant materials and growth condition

*Capsella rubella* materials used in the study were in *Cr22.5* ecotype background. The *Brassicaceae* species analyzed in this study were ordered from the germplasm bank of wild species of China (Extended Data Table 1). The seeds were germinated on MS medium 1% sucrose and 0.8% agar containing 10 µM Gibberellin (Sigma-Aldrich, no. G7645) at 22°C. The 10-day-old seedlings were then transplanted into soil in a controlled environment room (CER) at 22°C, 16 hrs light/8 hrs dark conditions.

### Plasmid construction and plant transformation

To generate the cell membrane marker line, the *pUBQ10:acyl-YFP* plasmid^1^ were transformed into the *Cr22.5* ecotype background. For the construction of the *pCrSTM:GUS* (Carub0002s0262, 1966 bp), *pCrWUS:GUS* (Carub0003s3208, 1982 bp), *pCrKNAT2* (Carub0002s1721, 4400 bp), *pCrKNAT6* (Carub0001s2296, 6074 bp), *pCrCYCB1;1* (Carub0007s02791, 460 bp) and *pCrCYCB1;2* (Carub0006s0503, 821 bp) GUS reporter plasmid, the respective promoter was isolated via PCR using Phusion High Fidelity Enzyme (New England Biolabs, no. M0530L) and inserted upstream of *GUS* gene of *pCambia*1301 vectors. For the *CrSTM* promoter activity analysis, a series of deletions were generated using the *pCrSTM:GUS* plasmid as the templet, the sequences were inserted upstream of *GUS* gene of *pCambia*1301 vectors. For generating the GFP-tagged transgenic lines, the full length genomic fragment plus the promoter sequences, *pCrSTM:CrSTM:GFP* (4986 bp), *pCrARF6:CrARF6:GFP* (Carub0001s2855, 7157 bp), *pCrARF8:CrARF8:GFP* (Carub0007s3229, 7151 bp), *pAtSTM (WT/mu):AtSTM:GFP* (AT1G62360, 4490 bp), were amplified and fused in-frame with the GFP coding sequence of *pCambia*1302 vector. For construction of the DEX-inducible *35S:CrSTM:GR* plasmid, the full length of *CrSTM* coding sequence was isolated from the fruit cDNA and inserted in-frame with GR in the pGTI0283 vector. For construction of two-component DEX-inducible *pLhGR>>amiR-CrSTM* plasmids, the artificial microRNA targeting the third exon of *CrSTM* were designed according to Ref 2. Briefly, the microRNA was synthesized as oligonucleotides and substitute the functional sites in the microRNA319 backbone via series PCRs. The resultant *amiR-CrSTM-miR319* PCR product was then integrated into the binary vectors using golden-gate cloning methods described in Ref 3. The *pLhGR>>CrSTM: 3ⅹFLAG* plasmid was generated in a similar way to *pLhGR>>amiR-CrSTM* except that *CrSTM* coding sequence was tagged with *3ⅹ* FLAG. For construction of the CRISPR/Cas9 genome editing plasmids, the DNA sequences encoding gRNAs adjacent to the PAM sequences (NGG) were designed using the CRISPR-P 2.0 software^4^ that targets first exon and eight sites targeting the promoter of *Carub0002s0262*, respectively. For gene editing the CrARF6/8, gRNAs targeting the exon 2 (*CrARF6*) and exon 1 (*CrARF8*), respectively, were designed. The gRNAs (Extended Data Table 2) were synthesized as oligonucleotides with golden-gate cloning adapters and were insert downstream of U6 promoters and then recombined with *pRPS5a:Cas9z:E9t* using golden-gate cloning methods to produce the binary vectors. All vectors were verified by sequencing and introduced into *Agrobacterium tumefaciens* strain LBA4404 by cold-shock transformation. The details of the primers were listed in the Extended Data Table 2.

Transformation of *Capsella* followed the floral dipping method optimized in Ref 3. The transformants were screened on MS plants with 1% sucrose and 0.8% agar containing either 40mg/L hygromycin (Roche, no. 10843555001) or 25mg/L DL-phosphinothricin (Duchefa, no. P0159.0250). For each construct, at least 10 independent transformants were obtained for further analysis. All the analysis were conducted on the T2 generation of the representative transgenic lines.

### Phenotyping and Scanning Electron Microscopy (SEM)

For whole-mount fruit phenotypes, stage 17 fruits of each genotype or Brassicaceae species were collected and photographed using Nikon D850 camera with a 105mm prime lens. For Scanning Electron Microscopy (SEM), the inflorescences of each genotype were fixed in FAA and then dissected in 70% ethanol. The samples were dehydrated and critically-point dried in CO_2_ following by spotter-coated with gold. The samples were examined using a Hitachi S-4800 scanning electron microscope (SEM) with an acceleration voltage of 10.0 kV.

### Protoplast preparation and scRNA-seq

The protoplasts from the fruit valve tips were prepared with the previous described protocol with minor modifications^5^. Briefly, the tip regions of stage-13 and stage-14 fruits were dissected from 200-260 fruits with a knife under a light microscope and then digested in the RNase-free enzymatic buffer (1.5% cellulase R10, 0.4% macerozyme R10, 0.4 M mannitol, 20 mM MES pH=5.7, 20 mM KCl, 10 mM CaCl2 and 0.1% BSA) for 2.5-3 h at 25 °C on a shaker at a speed of 40 rpm. The protoplasts were collected by cell strainers (40 μm in diameter, Falcon, no. 352340) and washed with W5 buffer (154 mM NaCl, 5 mM KCl, 125 Mm CaCl_2_ and 2 mM MES pH=5.7) 2-3 times to remove the cell debris and incomplete digested tissues on ice. The protoplasts were then concentrated with 0.6 M mannitol on ice followed by a quality test (determined by trypan blue staining as >90% viable cells to total cells for each sample). The concentration of protoplasts was finally adjusted to 1000-1200 cells/μL with 0.6 M mannitol. A total of 50000 cells were prepared for each sample. The scRNA-seq experiment were performed with two independent biological replicates for each stage.

To construct the sequencing library, the single-cell suspensions were loaded on a Chromium Single Cell Instrument (10× Genomics, Pleasanton, CA) to generate single-cell GEMs. Then, the cDNAs were synthesized by reverser transcription on a PCR machine. The sequencing library was generated using the Chromium Single Cell 3’ Gel Bead and Library Kit v3 (10× Genomics, no. 1000075 and 1000073) according to the users’ manufacture (Chromium Single Cell 3ʹ Reagent Kits v3, CG000183 Rev A). The libraries were sequenced by Illumina sequencer NovaSeq (Biomarker Technologies Beijing Co., Ltd) using two 150 bp paired-end kits. The raw scRNA-seq dataset was comprised of 28 bp Read1, 150 bp Read2 and 8 bp i7 index reads.

### Data integration, clustering and annotation

For aligning reads and generating gene-cell matrices, the raw dataset was processed with Cell Ranger 6.0.2 (10x Genomics) software with default parameters. The nuclear genome (version v1.1), the mitochondria genome and associated GTF annotation files of *Capsella rubella* were downloaded from Phytozome (https://phytozome-next.jgi.doe.gov/) and NCBI (https://www.ncbi.nlm.nih.gov/), respectively. These files were subsequently combined by the “cellranger mkref” function to build the reference. Then, the gene-cell matrix was generated by the “cellranger count” function. More than 90% reads in all the samples were mapped to the reference by executing the “cellranger count” function. To test the repeatability between samples, the “cellranger aggr” function was used to merge matrices of two replicates. All the detailed information of Cell Ranger is summarized in (Extended Data Table 3). One of two replicates (Mean Reads per Cell > 40000) from the replicates was used for further analysis.

The resultant gene-cell matrix was then processed into Seurat (v.4.3.0) package for in-depth data analysis including quality control, normalization, dimension reduction, clustering and annotation. For quality control, we applied the following criterions: (1) cells with unique molecular identifiers (UMIs) number 2500 ∼ 40000 were selected for analysis; (2) the percentage of mitochondrial UMIs was less than 5%; (3) cells containing expressed genes less than 200 were filtered out; (4) genes that were expressed in fewer than 3 cells were removed; (5) the genes with significant differential expression induced by the protoplasting process were excluded. To correct the variation caused by library preparation efficiency or sequencing depth, we used the “NormalizeData” to normalize the matrices. Top 2000 Highly Variable Genes (HVGs) were selected with the “FindVariableGenes” function by the vst method. For dimension reduction, the “RunPCA” function calculated 50 Principal Components (PCs) of HVGs and the top 20 PCs representing more than 85% accumulating contribution rate were selected for the downstream analysis. We next used the “FindNeighbors” function on the top 20 PCs to compute the nearest neighbor networks, then used the “FindClusters” function to cluster cells with “resolution = 0.7” argument based on Louvain method. The Uniform Manifold Approximation and Projection (UMAP) method was used to visualize cell clusters. The cluster specific or preferential expressed genes were identified by the “FindAllMarkers” function with “min.pct = 0.05, logfc.threshold = 0.5”. The “DotPlot” function and the “VlnPlot” function were used to define the enrichments of marker genes in the corresponding cell clusters. In the end, an additional R package “DoubletFinder” (v.2.0.2) was used to identify and remove the predicted doublet droplet from the clusters. The “plotly” (v.4.10.1) package was used to generate a 3D UMAP scatter graph.

### Differentiation trajectories analysis

For reconstructing the developmental trajectory of the epidermal cells, epidermal cells (Stage-13, cluster 4, 11, 12; Stage-14, cluster 3, 12, 14) were collected. To minimize the influence of stomata lineage on trajectory reconstruction, the cells expressing *FAMA* were removed. In order to normalize mRNA difference between cells, the “estimateSizeFactors” function and the “estimateDispersions” function were applied. Subsequently, monocle (v.2.18.0) package was used to infer the differentiation trajectory. A semi-supervised method was used to infer the genes involved in the corresponding biology process. The core developmental steps were calculated by the differential expression genes from the initial HVGs with “differentialGeneTest” function (screening criteria, qval < 0.01). We used the DDRTree algorithm with “reduceDimension” function to reduce the dimensions. To align cells in the trajectory, the “orderCells” function was performed, the root was established by the maximum expression of *ATML1*, *FDH*, *DCR* in trajectory. To visualize genes involved in cell cycle and cytokinesis (Extended Data Table S4), we used the “plot genes in pseudotime” function. The number of cells expressing these genes and the expression value of corresponding genes was calculated from counts and visualized using ggplot2 (v.3.4.2) package.

### Chemical treatment and phenotypic quantification

For the application of cyclin-dependent kinase inhibitor Flavopiridol hydrochloride (FVP), 100 μM working solutions were prepared with water and silwet (0.02%). The FVP and mock (water with 0.02% silwet) solutions were dipped onto the stage 13 and stage 14 fruits under the microscope. The plants were then kept in the CER under long-day condition (22°C, 16 hrs light/8 hrs dark). The fruit phenotype was recorded in 36 hours intervals. For dexamethasone (DEX, Sigma-Aldrich, no. D4902) and Cycloheximide (CHX, Sigma-Aldrich, no. 01810) treatment, 10 mM (DEX) and 2 mM (CHX) stock solutions were resolved in Dimethyl sulfoxide (DMSO, Sigma-Aldrich, no. D8418). For the *35S:CrSTM:GR* lines, 10-days old seedlings were transferred onto the MS medium containing either 10 μM DEX, 2 μM CHX, 10 μM DEX plus 2 μM CHX or mock (equal amount of DMSO) for 4 hours on a shaker in the CER at a speed of 40 rpm, after which the samples were harvested in liquid nitrogen and subjected to gene expression analysis. For the *pLhGR>>amiR-CrSTM* lines, 20 μM DEX solutions with 0.02% silwet or mock (water with 0.02% silwet) solutions were dipped onto the inflorescence and kept in a humid condition for 24 hour, then the plants were kept in the growth chamber under long-day condition (22°C, 16 hrs light/8 hrs dark). The fruit phenotype was recorded in the respective developmental stages with Nikon D850 camera or SEM. For the *pCrSTM (1.5 kb WT/mu):GUS/pLhGR>>CrSTM:3ⅹFLAG* lines, 10-days old seedlings were transferred into the liquid MS medium containing either 10 μM DEX or mock (equal amount of DMSO) for 12 hours on a shaker in the CER at a speed of 40 rpm. The samples were subjected to GUS staining at 4 hours intervals.

The fruit shape was quantified as the shoulder index value, which was calculated with the anti-trigonometric function θ=Arctan((L1-L2)/W) using the parameters described in Ref 6.

### Live imaging and cell behavior analysis

To conduct live imaging experiment, the *pUBQ10:acyl-YFP* plants were cultivated on soil in the CER under long-day conditions until they reached the bolting stage (22°C, 16 hours light/8 hours dark). At stage 12, the fruits were dissected, transferred onto a microscope slide and imaged capturing YFP signals at 24-hour intervals using a Zeiss inverted laser confocal microscope (Zeiss LSM 980) equipped with a water immersion objective (x25/0.95). The excitation wavelength was set at 513 nm, and the emission window approx. 530 nm. Confocal stacks were acquired at a resolution of 1024×1024, with a distance of less than 0.5 μm in the Z-dimension. During the intervals between imaging sessions, the samples were maintained on Petri dishes containing 1/2 MS medium supplemented with vitamins (Coolaber, no. PM1011) and 1% sucrose in the CER under long-day conditions (22°C, 16 hours light/8 hours dark). The acquired images were analyzed using MorphoGraphX^7^. To quantify cell area ratio, growth anisotropy and proliferation, the fluorescence was projected into a mesh, cell outlines were segmented, and the relationships between cells were indicated across successive time points. Heatmaps illustrating the differences between two time points were generated, with the heatmap being shown on the fruit stage at the latter stage. Representative growth or proliferation maps were obtained from a single experiment.

### RNA extraction and expression analysis

The samples (fruits or seedlings) subjected to expression analysis were immediately fixed in liquid nitrogen. Total RNA was isolated from the samples using the SV Total RNA Isolation System, and DNase I was added to digest the genomic DNA (Promega, no. Z3100) following the manufacturer’s instructions. 500ng of total RNA was reverse transcribed into Complimentary DNA (cDNA) in a 10μl reaction with the SuperScript™ IV First-Strand Synthesis System (ThermoFisher, no. K1622) according to the manufacturer’s instructions.

For real-time qPCR, gene specific primers were designed (Extended Data Table 2) and verified by PCR followed by sequencing. The SYBR Premix Ex Taq (TaKaRa, no. RR420DS) was used to perform real-time qPCR with ROX as a reference dye on a StepOne Plus Real-Time PCR System (Life Technology). The CT value of each gene was determined by normalizing the fluorescence threshold. The relative expression level of the target gene was determined using the ratio = 2^-ΔCT^ method, and *CrUBQ10* was used as an internal control.

GUS histochemical assays were performed as previously described^6^.

### Chromatin immunoprecipitation (ChIP) and ATAC-Seq

For the ChIP experiment, stage12-15 fruits from each genotype (GFP-tagged transgenic lines, see “Plasmid construction and plant transformation” section) and WT plants were harvested and cross-linked in 1 x PBS plus 1% formaldehyde under vacuum for 10 mins then add 2 M glycine to a final concentration of 125 mM and apply vacuum for another 5 mins. Each sample containing ~3.0 g of tissue was ground in liquid nitrogen into fine powder and nuclear was isolated by Honda buffer (0.44 M sucrose, 1.25% Ficoll, 2.5% Dextran T40, 20 mM Hepes KOH pH=7.4, 0.5% Triton X-100, 10 mM MgCl_2_ and 2 tablets Protease Inhibitor Cocktail/100ml), chromatin was released and fragmentated by sonication. After sonication, ~1/12 (50 μL) DNA sample was taken out as Input. The remaining DNA was subjected to anti-GFP immunoprecipitation using Pierce Protein G Magnetic Beads (ThermoFisher, no. 88847) coated with monoclonal anti-GFP antibody (Roche, no. 11814460001) at 4°C for 4 hrs. After the immunoprecipitation, beads were then washed with the immunoprecipitation buffer and TE buffer then processed into the reverse crosslinking procedural in presence of 10% SDS at 65°C for 12 hours. The proteins in the DNA complex were digested by Proteinase K treatment at 45°C for 1 hour. After phenol/chloroform extraction, DNA was visualized by Glyco-Blue (ThermoFisher, no. AM9515) and precipitated in 70% ethanol, dried and resolved in ChIP-grade water (Sigma-Aldrich, no. W4502). qPCR was performed using SYBR Premix Ex Taq on a StepOne Plus Real-Time PCR System (Life Technology).

The ATAC-seq experiment was performed following the protocol as previously described^8^ with minor modifications. Briefly, ~0.5 g of the stage 12-14 and stage 15-16 fruit were harvested and were ground into fine power in liquid nitrogen, respectively. Approximately 50,000 nuclei were collected and washed using cold PBS, and then resuspended with cold lysis buffer (10 mM Tris-HCl, pH 7.4, 10 mM NaCl, 3 mM MgCl_2_, 0.1% IGEPALCA-630). The nuclei were then subjected to transposing reaction with Tn5 Transposase at 37 °C for 30 min, and the DNA was purified using a Qiagen MinElute PCR Purification Kit (Qiagen, no. 28006). The sequencing libraries were prepared by PCR amplification using the NEBNext PCR master mix (New England Biolabs, no. M0541S). The amplified libraries were purified with AMPure beads (Beckman Coulter, no. A63880) and sequenced on the Illumina platform to generate 150-bp, paired-end reads. For the ATAC-seq data analysis, the raw sequencing reads were first trimmed by fastp (v.0.20.0) to generate the clean fastq files. After filtering out the low-quality reads (reads length below 35 and bases quality value Q less than 10), the sequences were aligned to the *Capsella rubella* reference genome^9^ using Bowtie2 software^10^. The duplicated reads were marked and removed by sambamba (v.0.6.7) and bedtools (v.2.25.0)^11^. Peaks were then called using MACS2 (v2.1.4)^12^ software with a screening criteria of false detection rate (FDR) < 0.05. Common peaks present in both replicates were identified and used for the following-up analysis. DeepTools (v.3.1.2) was used to map the density distribution of sequencing read in the upstream and downstream of the Transcription Start Site (TSS) of each gene. For visualization, the datasets were converted to bigwig formats using BamCoverage using DeepTools (v.3.1.2) with a bin size of 10 bp and then visualized using the Integrative Genomics Viewer (IGV, v.2.4.14) software. The ATAC-seq experiments were performed with two independent biological replicates.

### Evolution analysis of the STM-binding sites

The promoter sequences of STM-orthologs were isolated by TAIL-PCR (TAKARA, no.6108) from the DNA of the representative Brassicaceae species (Extended Data Table 1). The 20-bp orthologous region containing the STM-binding site in the centre was extracted and aligned using MUSCLE v3.8^13^ with default settings. For the sample panel, we divided them into two groups, one containing the species with the STM-binding site (*n* = 21), the other containing the species without STM-binding site (*n* = 28). These two aligned files were separately inputted into the WebLogo online tool (https://weblogo.berkeley.edu/) to generate the sequence logo.

To test whether the 6-bp STM-binding element is under strong selective constraint, we conducted a comparison of the 6-bp STM binding site from the aforementioned two groups with neutral sites. We measured the conservation of every single site using the Shannon entropy of the nucleotide distribution of each column in the alignment. For the control, we use the coding sequences of STM-orthologous genes from Brassicaceae species with genome sequences in Brassicaceae, and then extracted all the four-fold degenerate sites (4-fold sites) using a script (extract_4fold_degenerate_sites.py from https://github.com/mscharmann/tools/blob). We next randomly sampled 6 sites from all the 4-fold sites each time, calculated the mean of Shannon entropy values, and repeated these two steps for 1000 times. Statistical tests among the three groups were performed using two-sided Mann-Whiteney U test in R (v.4.1.2).

### Quantification and statistical analysis

All statistics were calculated in Microsoft Excel. All measured data are presented as means ±SD specified along with sample sizes (n) in the methods and in figure legends. Comparisons between groups for the analysis of qRT-PCR, fruit characters and ChIP-qPCR was performed with Microsoft Excel student’s *t*-test, and significance levels are marked as: ∗ *p* < 0.05, ∗∗ *p* < 0.01, ∗∗ *p* < 0.001.

## Acknowledgements

We are grateful to Yu-Ling Jiao for the fruitful discussion on the preliminary data and to Mark Bal for critically reading the manuscript and providing comments prior to submission. We thank Phil Robinson for photographic assistance and Dr. Nobutaka Mitsuda for providing the *35S:MicroRNA319* plasmid. This study was supported by a grant from the Biotechnological and Biological Sciences Research Council (BBSRC) (BB/P020747/1 to L.Ø.), a BBSRC Institute Strategic Programme Grant to the John Innes Centre (BB/P013511/1), the National Key Research and Development Program of China (2022YFF1301704 to Ya.D.), grants from the National Natural Science Foundation of China (32170227 and 32221001 to Ya.D.) and K.C. Wong Education Foundation (GJTD-2020-05, received by Ya.D.).

## Author contributions

Ya.D. initiated, conceived, and designed the research with support from L.Ø. Ya.D. and L.Ø. supervised the project. Ya.D. performed all the research with assistance from Z.H., M.M., H.S., and Y.Z. Z.H., Z.H., Yi. D. and T.S. were involved in the promoter analysis. M.M. and R.S.S conducted the cell growth analysis. H.S. performed the scRNA-seq data analysis. F.G. collected the Brassicaceae species and extracted the DNA. Y.Z., Q.Y., and T.L. carried out phylogenetic analysis, fruit morphological analysis and gynoecia SEMs. G.X. did the molecular evolution analysis of STM-binding site. Ya.D. and L.Ø. outlined and wrote the manuscript. All authors participated in the discussion of the data and in the production of the final version of the manuscript.

## Extended Data

**Extended Data Fig. 1.**
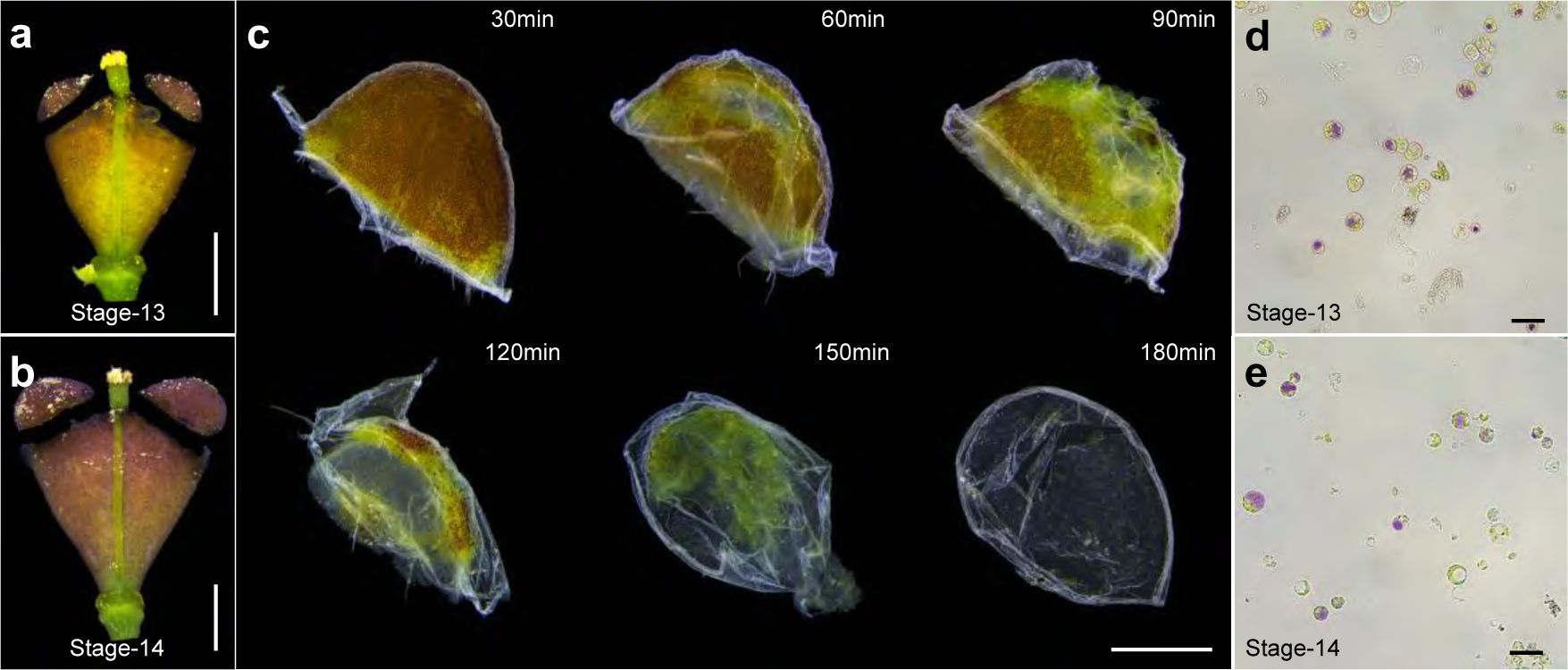
Sample preparation for the scRNA-seq analysis. **a** and **b**, Sampling stages. The valve tips of stage-13 (**a**) and stage-14 (**b**) fruit were harvested for protoplast preparation. **c**, Time-course record of the enzymatic digestion process of the stage-14 valve tips. Please note that the digestion was nearly complete after 150 min and fully complete after 180 min. **d** and **e**, The final protoplasts used for scRNA-seq analysis. The protoplasts from stage 13 (**d**) and stage 14 (**e**) fruit were shown. Scale bars, **a** and **b**, 300 μm; **c**, 150 μm; **d** and **e**, 50 μm.

**Extended Data Fig. 2.**
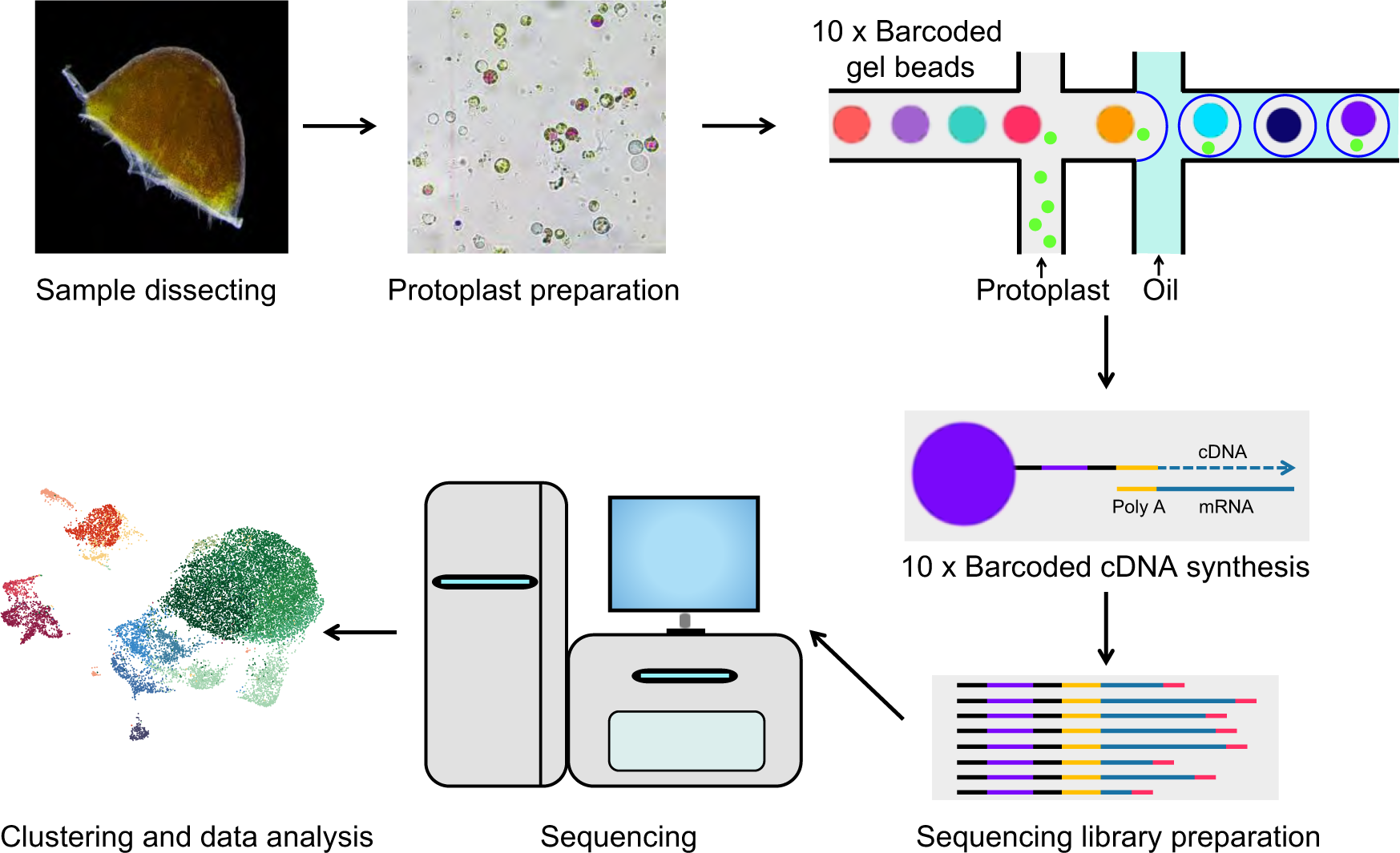
Workflow of the scRNA-seq analysis process. The fruit valve tips were dissected from fruit and digested to protoplast using RNase-free enzymatic buffer. The single-cell suspensions were loaded on a 10x platform to generate single-cell GEMs. After library preparation and sequencing, raw data was counted using Cell Ranger software to get a gene-cell matrix, and the Seurat package was used to visualize a UMAP plot for sample scRNA-seq transcriptome.

**Extended Data Fig. 3.**
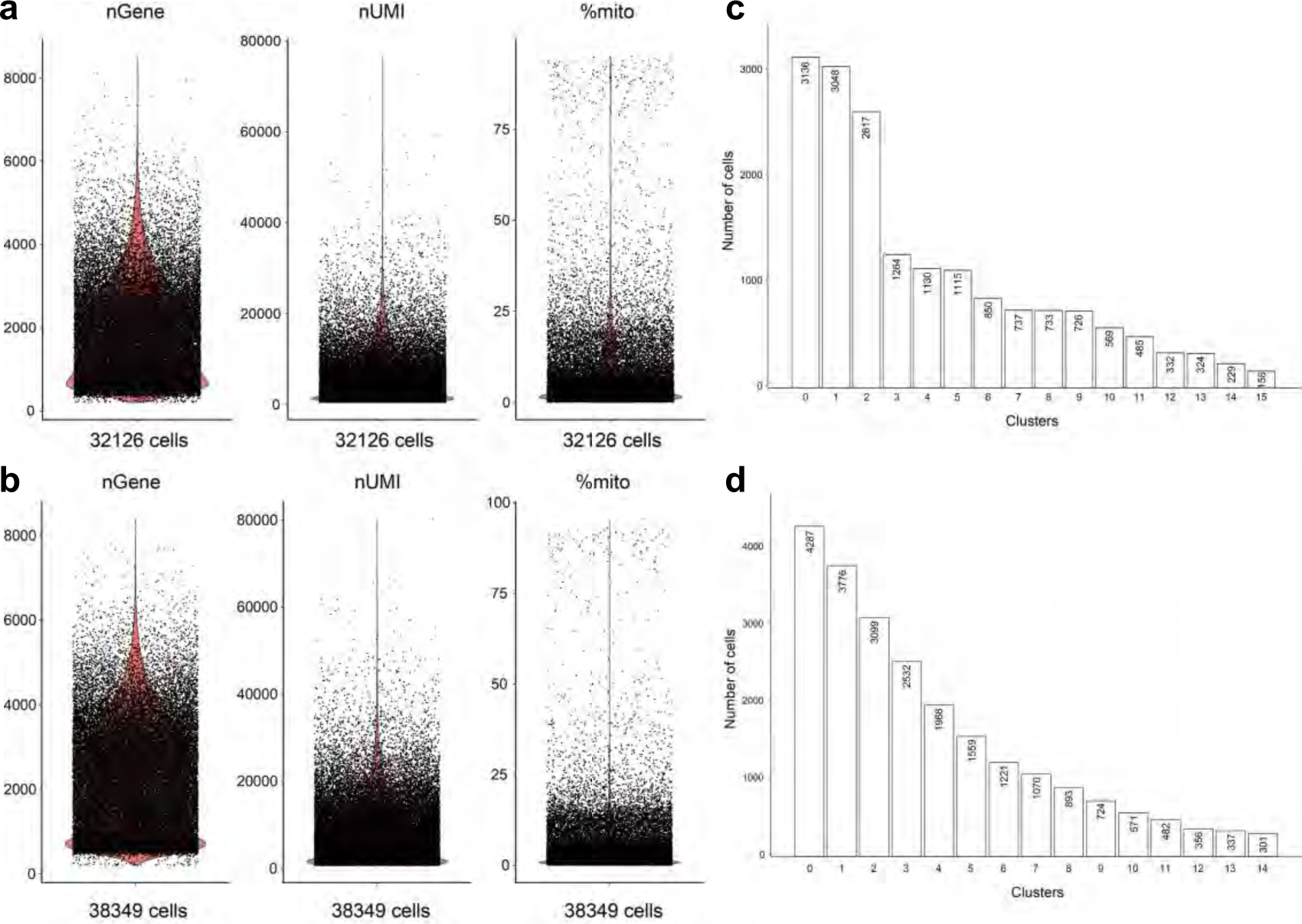
scRNA-seq data quality parameters. **a** and **b**, The basic parameters of scRNA-seq data of stage 13 (**a**) and stage 14 (**b**) fruit valve tips. nGene indicates the number of unique genes (left panel), nUMIs indicates the number of unique molecular identifiers (middle panel) and %mito indicates the percentage of mitochondrial genes expressed in the total scRNA-seq transcriptome (right panel). We captured 32126 cells for stage 13 fruit samples and 38349 cells from stage 14 fruit samples, respectively. **c** and **d**, The cell numbers in each cluster from stage 13 fruit samples (**c**) and stage 13 fruit samples (**d**).

**Extended Data Fig. 4.**
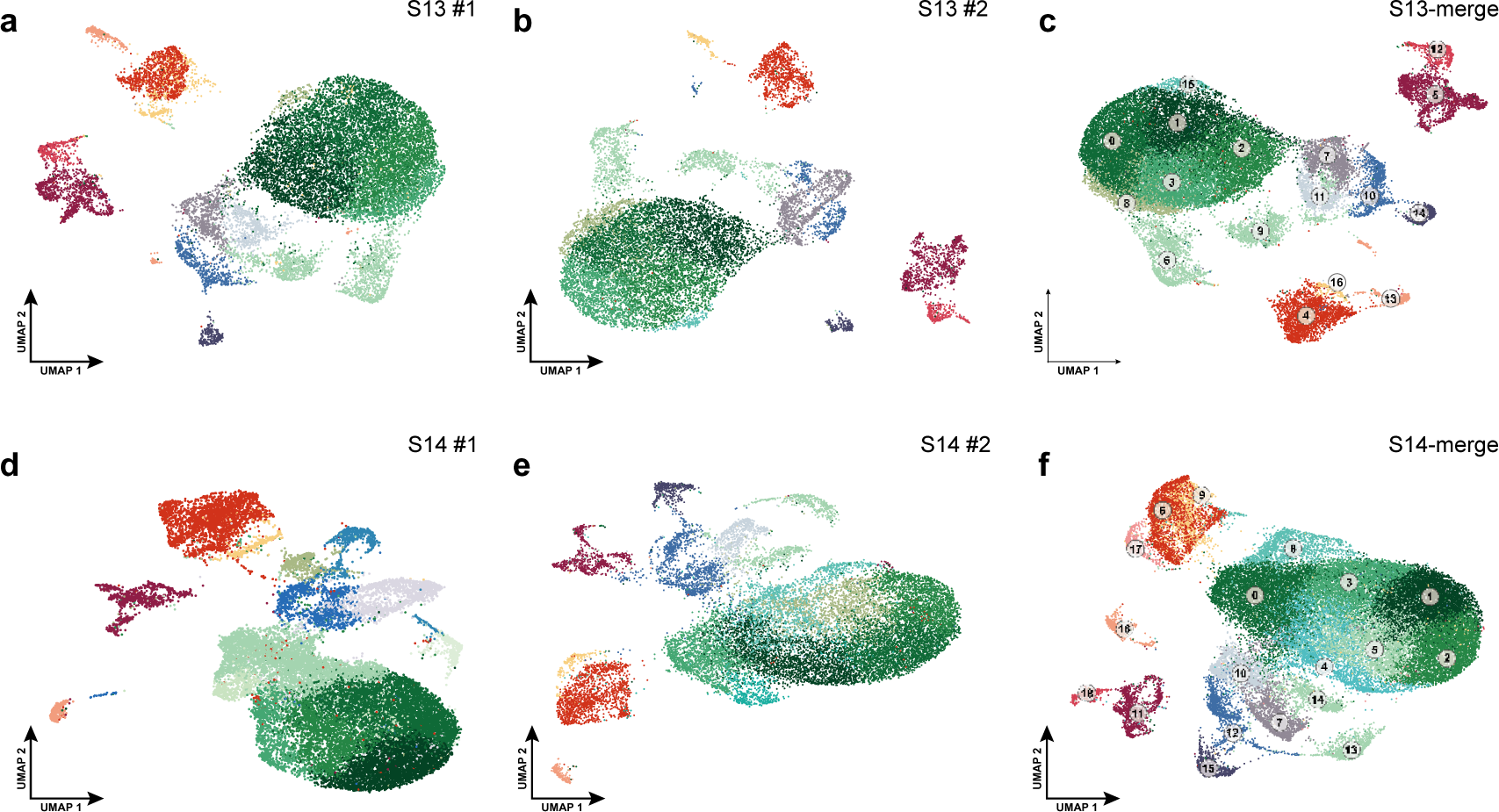
Reproducibility test of scRNA-seq data. **a**, UMAP projection of scRNA-seq datasets replicate 1 of stage 13 (S13) fruit sample. **b**, UMAP projection of scRNA-seq datasets replicate 2 of stage 13 fruit sample. **c**, UMAP projection of scRNA-seq datasets combined by replicate 1 and replicate 2 from stage 13 fruit sample. **d**, UMAP projection of scRNA-seq datasets replicate 1 of stage 14 (S14) fruit sample. **e**, UMAP projection of scRNA-seq datasets replicate 2 of stage 14 fruit sample. **f**, UMAP projection of scRNA-seq datasets combined by replicate 1 and replicate 2 from stage 14 fruit sample. Please note that UMAP projection of a single replicate or combined data set produces highly similar cell clusters, indicating high reproducibility of the experiment. Cells and clusters are annotated as shown in Fig. 2a and 2b.

**Extended Data Fig. 5.**
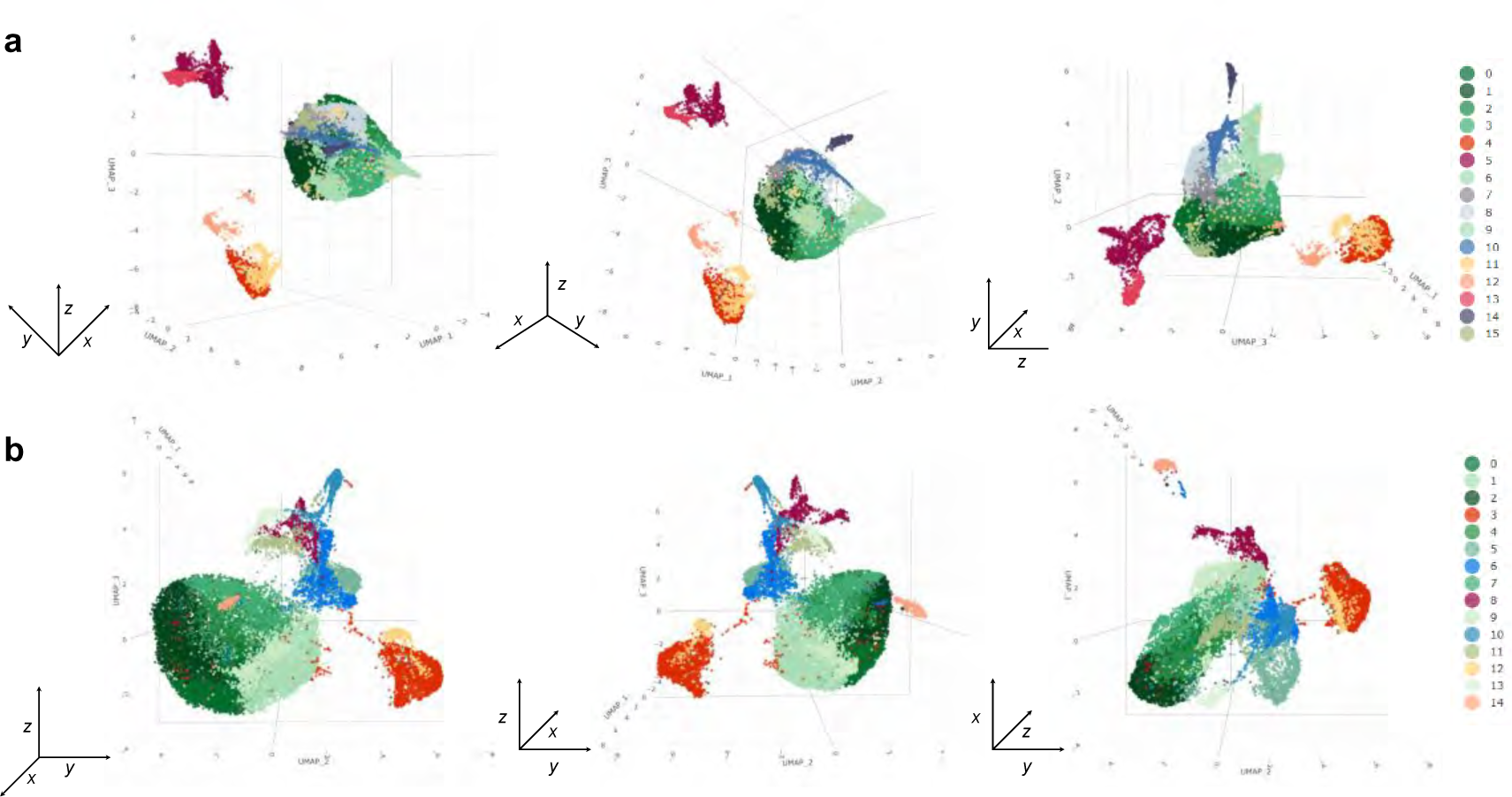
3D views of the scRNA-seq transcriptomic cell clusters. **a**, Visualization of cell clusters by 3D UMAP scatterplots of stage 13 fruit samples. **b,** Visualization of cell clusters by 3D UMAP scatterplots of stage 13 fruit samples. Cluster name and colors are the same as in Fig. 2a and 2b.

**Extended Data Fig. 6.**
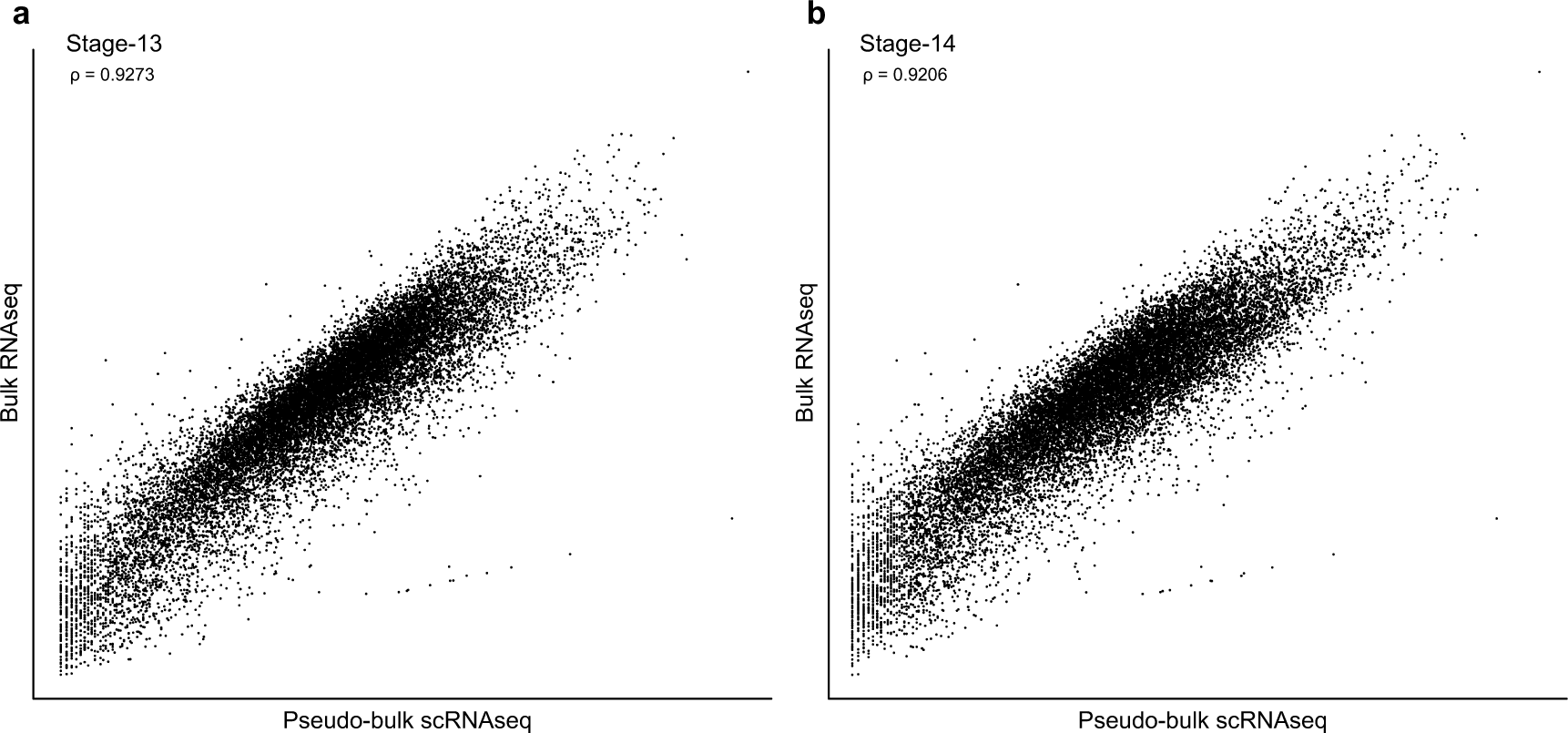
Correlation analysis of scRNA-seq and bulk RNA-seq data. **a**, The correlation index of fresh tissue bulk RNA-seq data and pseudo-bulk scRNA-seq data of stage 13 fruit sample. **b**, The correlation index of fresh tissue bulk RNA-seq data and pseudo-bulk scRNA-seq data of stage 13 fruit sample. The indexes are calculated by Spearman’s rank correlation method.

**Extended Data Fig. 7.**
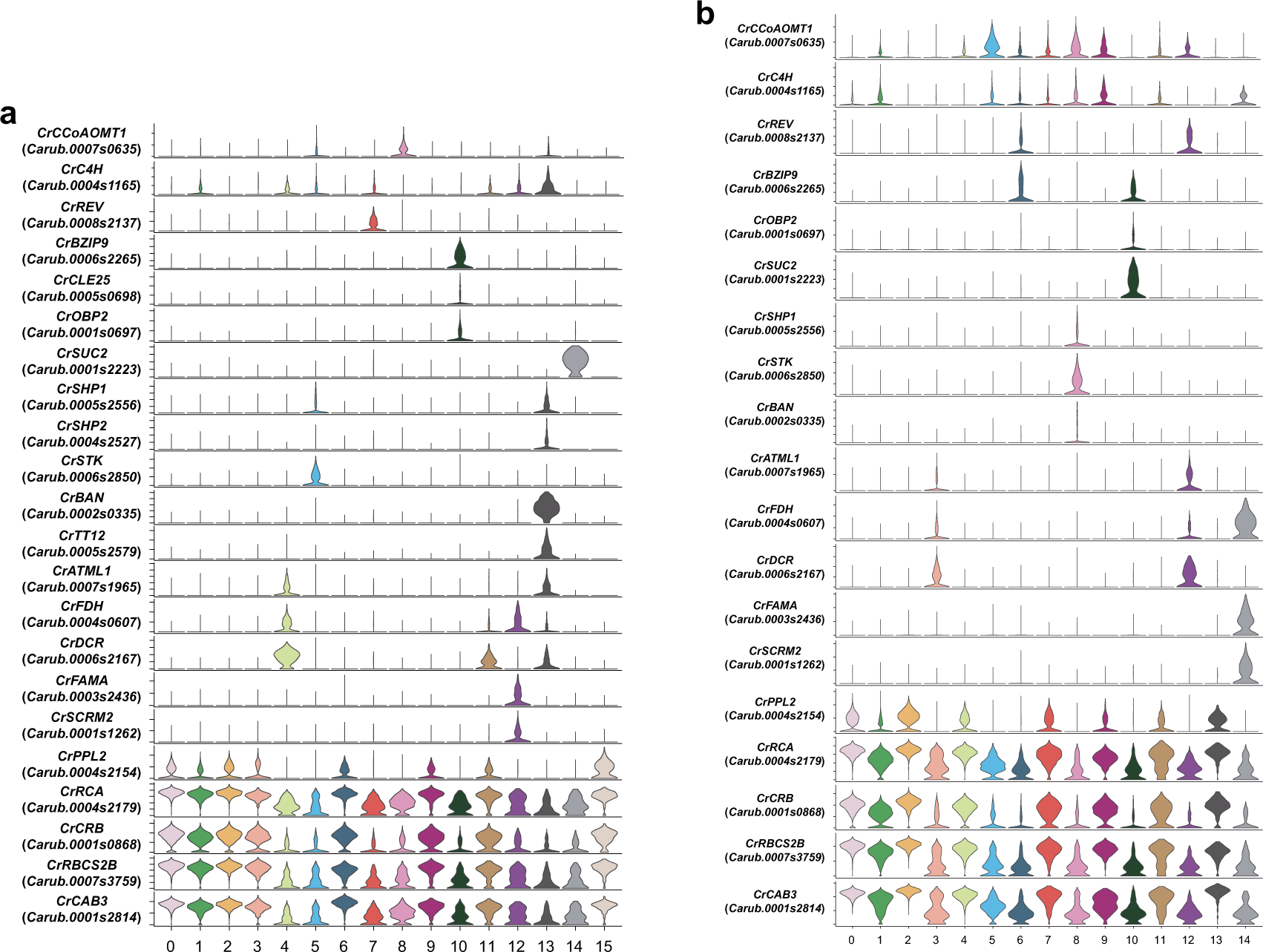
Expression patterns of representative cluster-specific makers genes on UMAP. **a**, Expression patterns of markers genes in stage 13 cell altas by Violin plot. **b**, Expression patterns of markers genes in stage 14 cell altas by Violin plot. *CrCCoAOMT1* and *CrC4H* are used to identify endocarp cell clusters. Vascular cell clusters are classified by the expression of *CrREV*, *CrBZIP9*, *CrCLE25*, *CrOBP2* and *CrSUC2*. Embryo cell clusters are identified by expressing transcript of *CrSHP1*, *CrSHP2*, *CrSTK*, *CrBAN* and *CrTT12*. *CrATML1*, *CrFDH* and *CrDCR* are related to the epidermis cell clusters. *CrFAMA* and *CrSCRM2* are specifically expressed in guard cell clusters. Mesophyll cell clusters were identified by photosynthetic related genes, such as *CrPPL2*, *CrRCA*, *CrCRB*, *CrRBCS2B* and *CrCAB3*. The full name of selected genes and associated references are provided in Extended Data Sheet 2.

**Extended Data Fig. 8.**
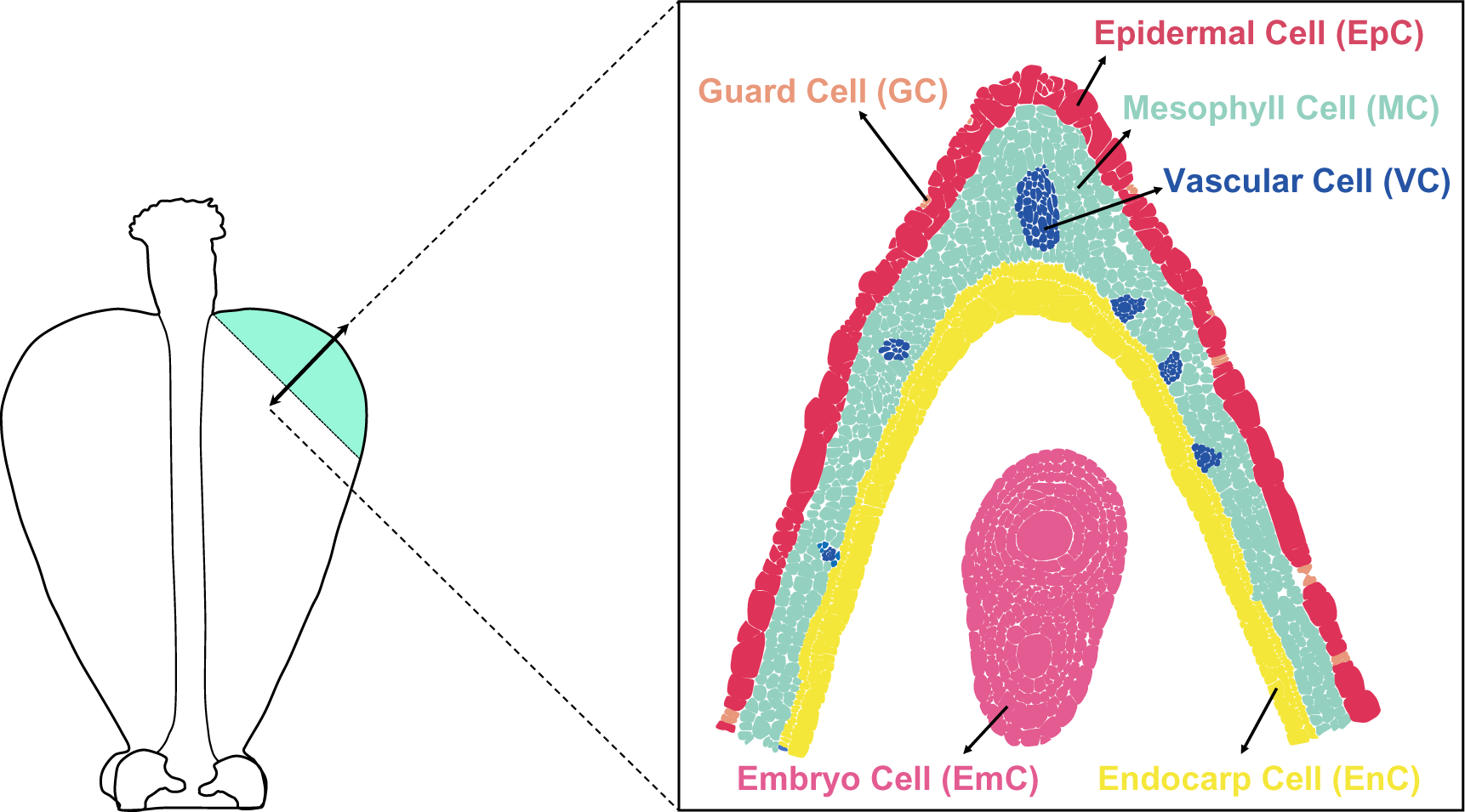
Schematic of anatomy and cell types of the valve tips. The double-head arrow indicates the direction of the section from a stage-13 fruit. Note that different cell types or tissues were colored based on the section in a 1:1 ratio.

**Extended Data Fig. 9.**
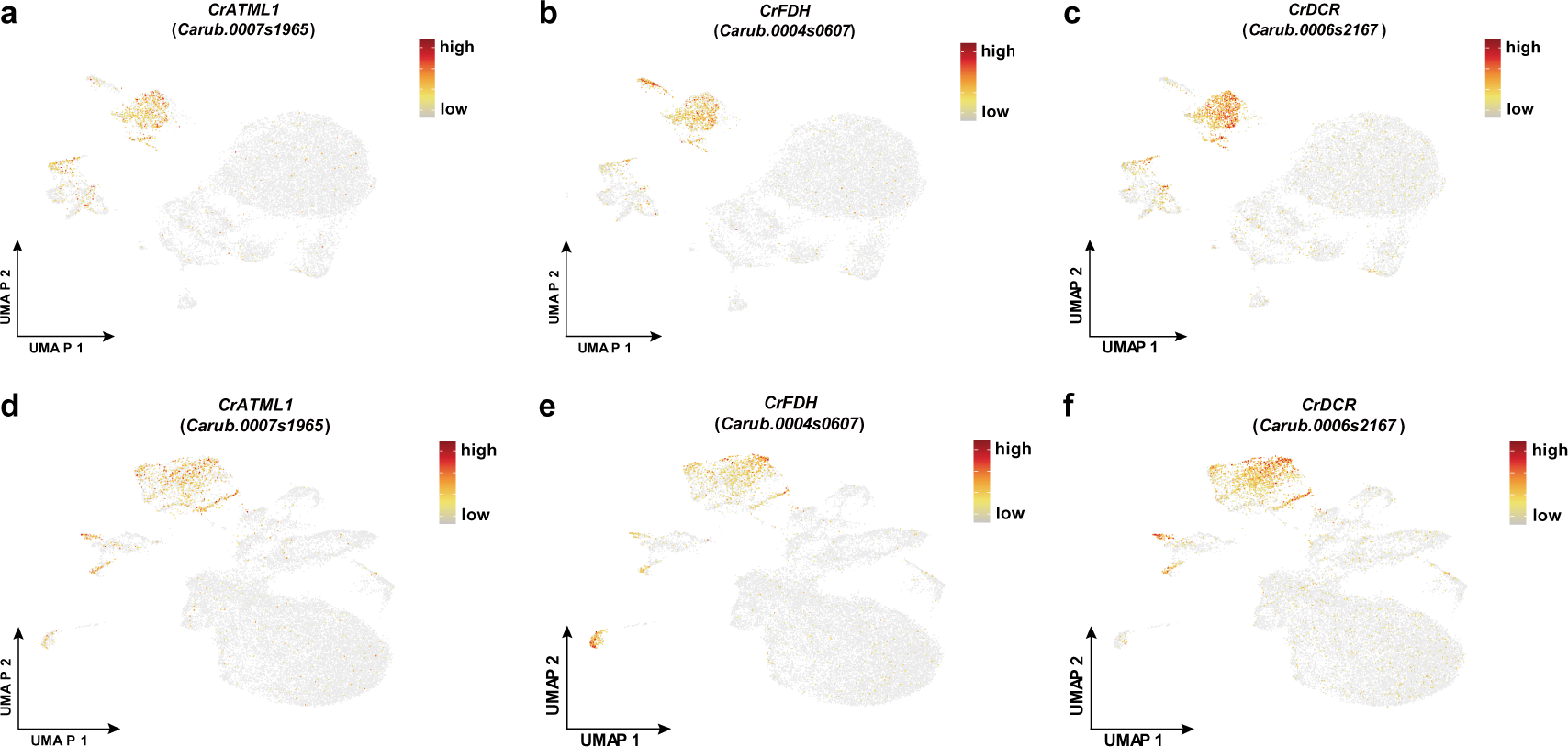
Expression of epidermal cell marker genes. **a**-**c**, UMAP plot showing the selected top markers genes expressed in the epidermal cells from stage 13 valve tips. **d**-**f**, UMAP plot showing the selected top markers genes expressed in the epidermal cells from stage 14 valve tips. **a** and **d**, *CrATML1*; **b** and **e**, *CrFDH*; **c** and **f**, *CrDCR*. The full name and referenced expression pattern of the selected genes are given in Extended Data Sheet 2.

**Extended Data Fig. 10.**
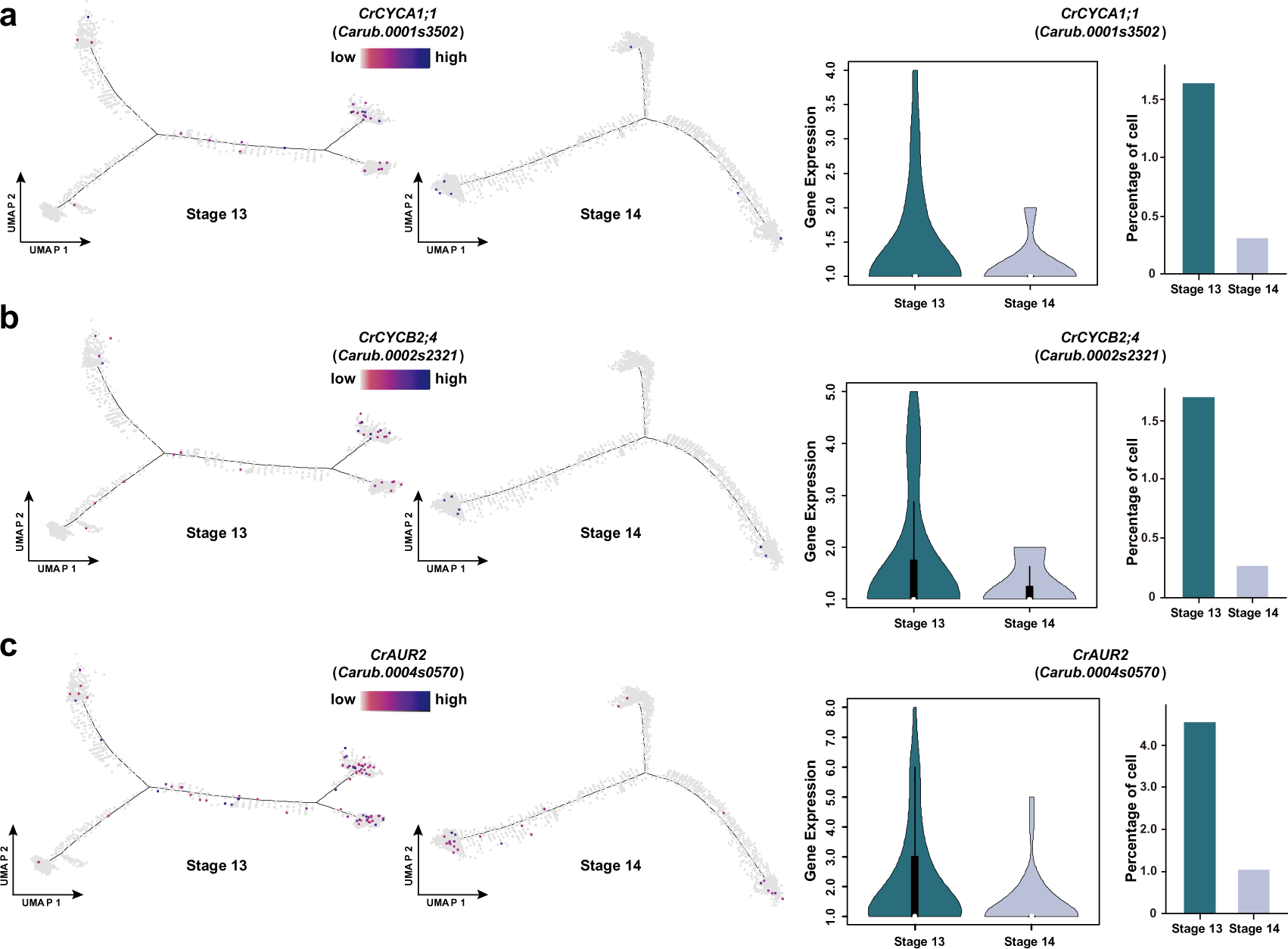
Gene expression profiles in the pseudotime differentiation trajectories. **a**-**c**, Comparisons of the expression pattern, expression level and the percentage of cells expressing the genes involved in cell cycle and cytokinesis (**a**, *CrCYCA1;1*; **b**, *CrCYCB2;4*; **c**, *CrAUR2*) in the valve tip epidermal cells between stage 13 and stage 14 fruits. The gene expression level is calculated by the ggboxplot function (R ggpubr package). Each dot in the differentiation trajectory indicates a single cell.

**Extended Data Fig. 11.**
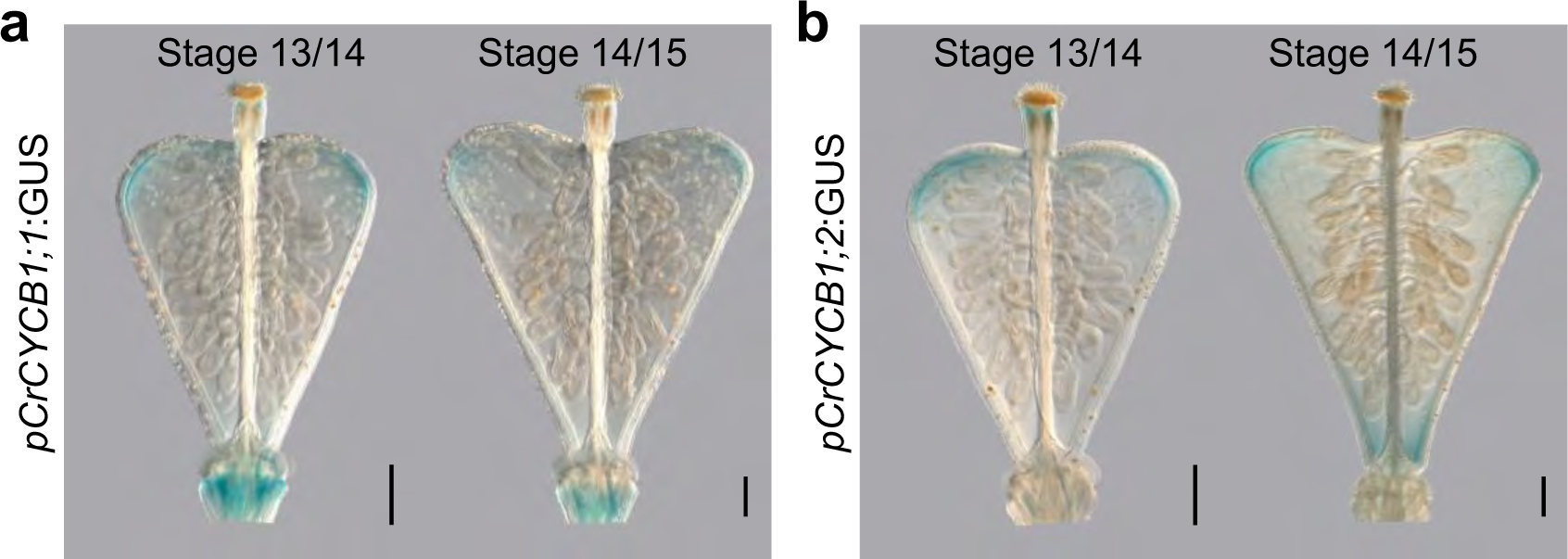
Expression analysis of two cyclin genes in the fruits. **a** and **b**, Expression pattern of *CrCYCB1;1* (**a**) and *CrCYCB1;2* (**b**) shown by GUS staining with respective reporter lines at developmental stages 13/14 and 14/15, showing a gradual decrease in the expression of these cyclin genes in the valve tips. Scale bars, 100 μm.

**Extended Data Fig. 12.**
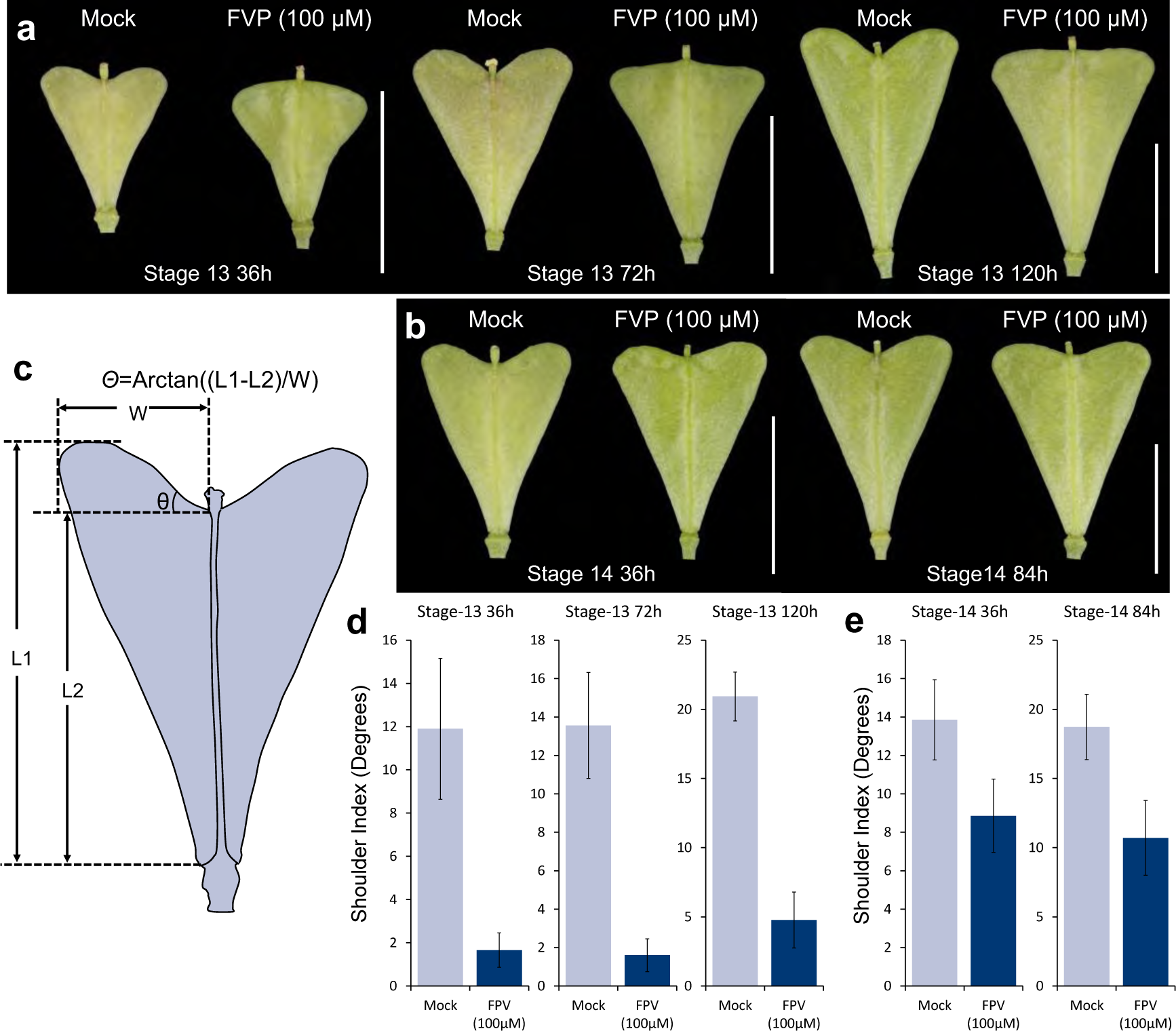
Effects of FVP treatment on *Capsella* fruit shape development. **a**, Time-course pictures of the fruits upon FVP and Mock treatment started from stage 13. Please note after 36 h, the stage-13 fruit has already developed to stage 14, 72 h to stage 15 and 120 h to stage-16. **b**, Time-course pictures of the fruits upon FVP and Mock treatment started from stage 14. Equivalent time points to stage-13 were shown. **c**, Schematic drawing to illustrate the shoulder index calculation^6^. **d**, Shoulder index measurements of fruits from FVP and Mock treatment started from stage-13. **e**, Shoulder index measurements of fruits from FVP and Mock treatment started from stage-14. Scale bars, 5 mm in **a** and **b**.

**Extended Data Fig. 13.**
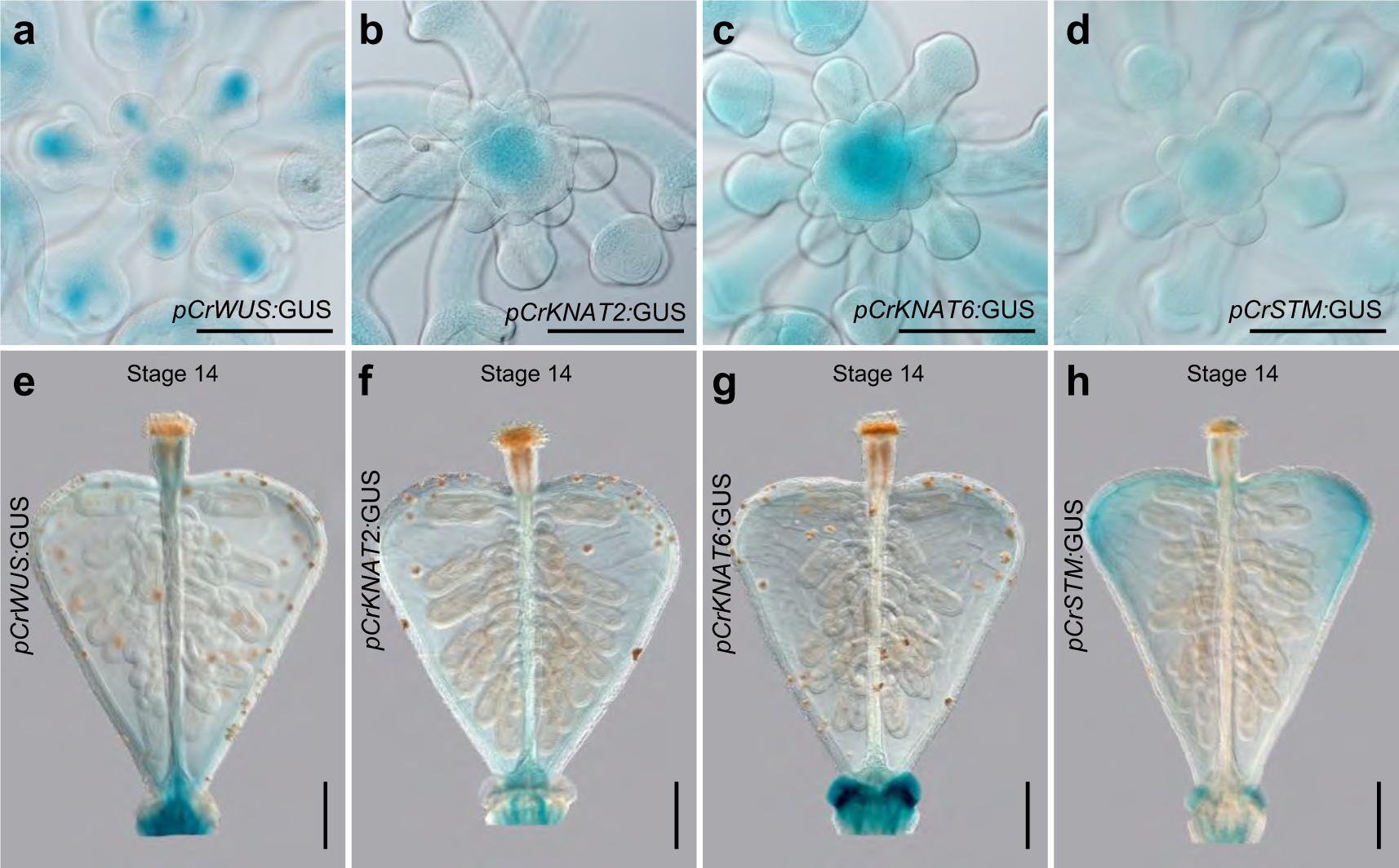
Expression analysis of genes involved in SAM development. **a**-**d**, Expression pattern of *CrWUS* (**a**) and *CrKNAT2* (**b**), *CrKNAT6* (**c**) and *CrSTM* (**d**) shown by GUS staining with respective reporter lines in the shoot apical meristem (SAM), showing all genes are expressed in the SAM. **e**-**h**, Expression pattern of *CrWUS* (**e**) and *CrKNAT2* (**f**), *CrKNAT6* (**g**) and *CrSTM* (**h**) shown by GUS staining with respective reporter lines in the stage-14 fruits, showing only *CrSTM* (**h**) expression was detected in the fruit valve tips. Scale bars, 100 μm.

**Extended Data Fig. 14.**
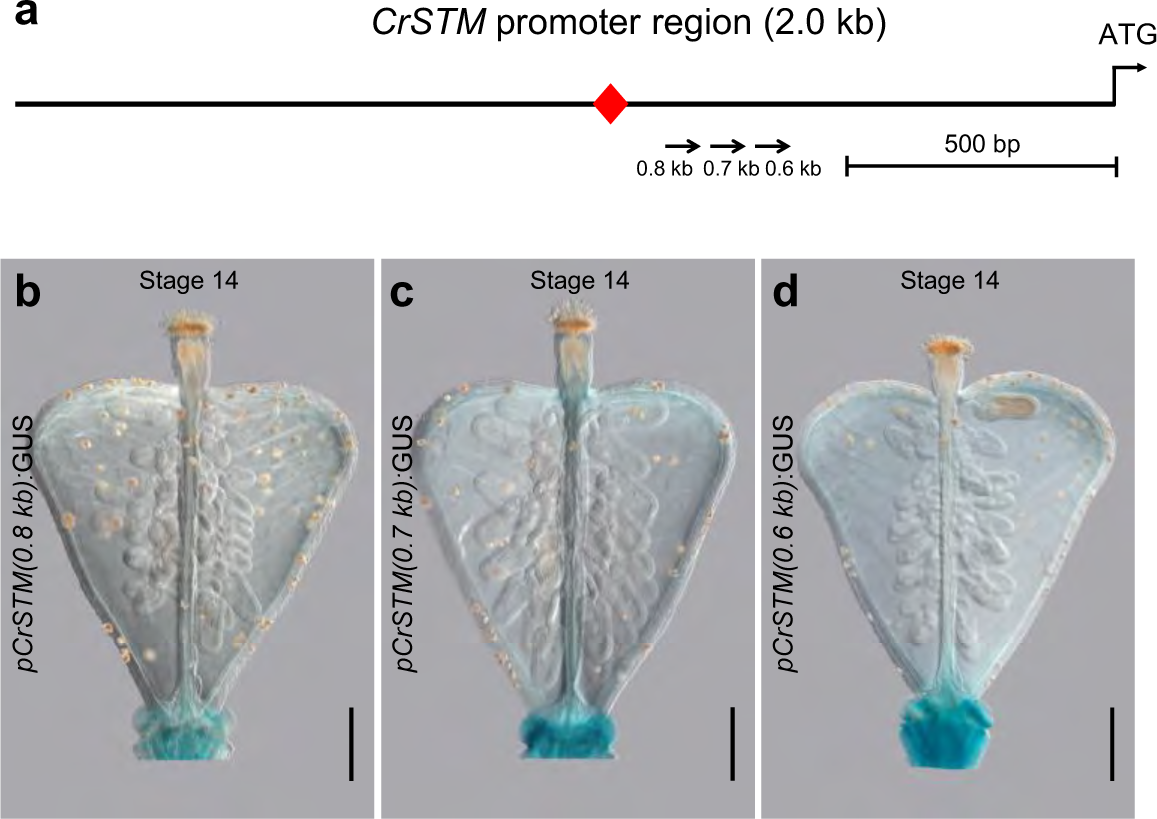
*CrSTM* promoter activity analysis by GUS staining. **a**, Schematic show of the 2.0 kb *CrSTM* promoter upstream of the start codon. The arrows indicate the promoter deletion series in the GUS reporter analysis. **b**-**d**, Expression pattern of *CrSTM (0.8 kb)* (**b**), *CrSTM (0.7 kb)* (**c**) and *CrSTM (0.6 kb)* (**d**) shown by GUS staining with respective reporter lines in the stage-14 fruits, showing no *CrSTM* expression could be detected in the fruit valve tips. Scale bars, 100 μm.

**Extended Data Fig. 15.**
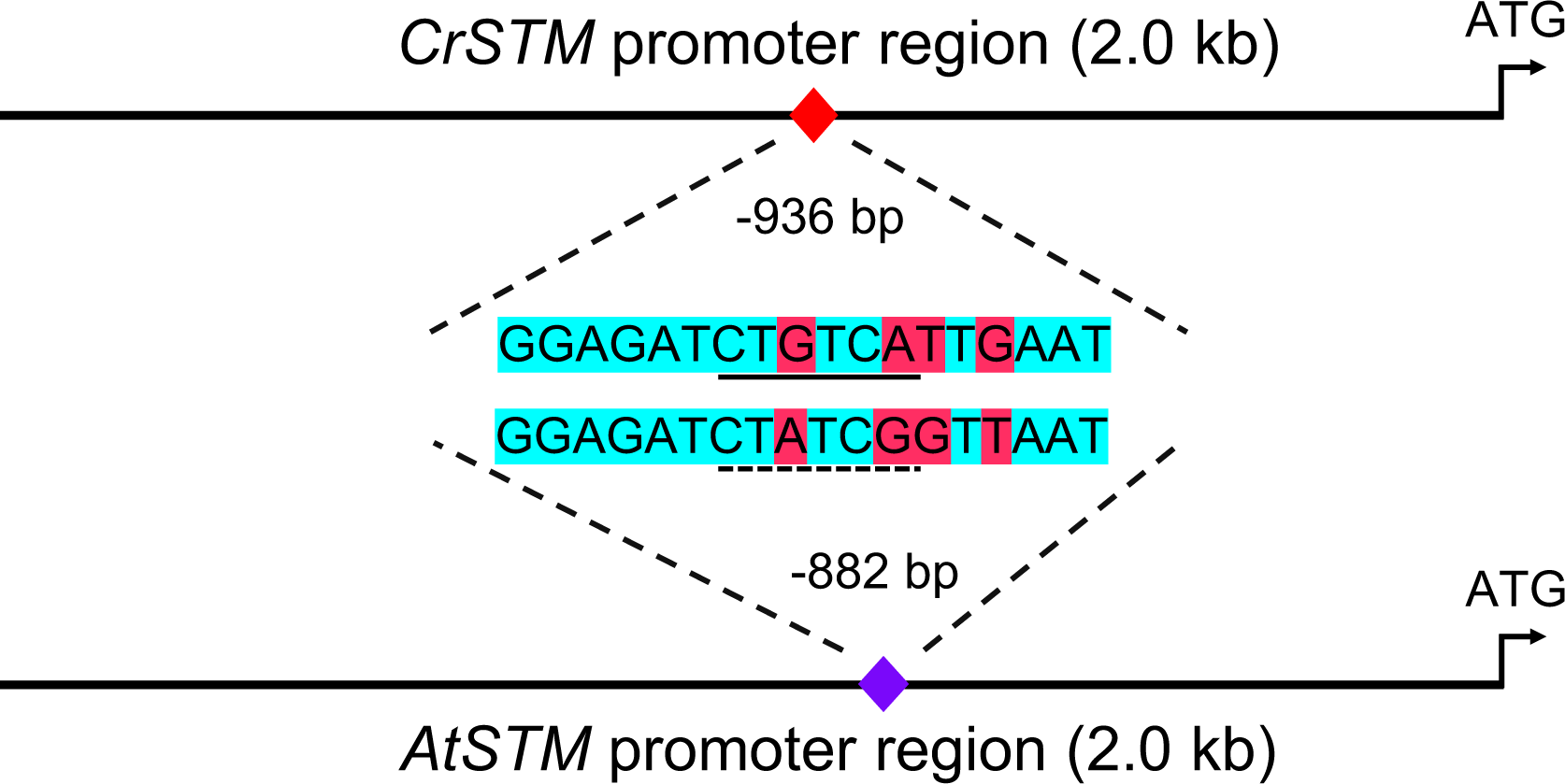
Sequence comparison of the *STM* promoters between *Capsella* and *Arabidopsis*. 2000 bp sequence upstream of the transcription start site (TSS) was aligned between *Capsella* and *Arabidopsis*. The STM-binding site (underlined) is located at -936 bp position in the *CrSTM* promoter, which is homologous to the -882 bp region in the *AtSTM* promoters. The identical sequences were shown in green and differentiated sequences were shaded in red, respectively.

**Extended Data Fig. 16.**
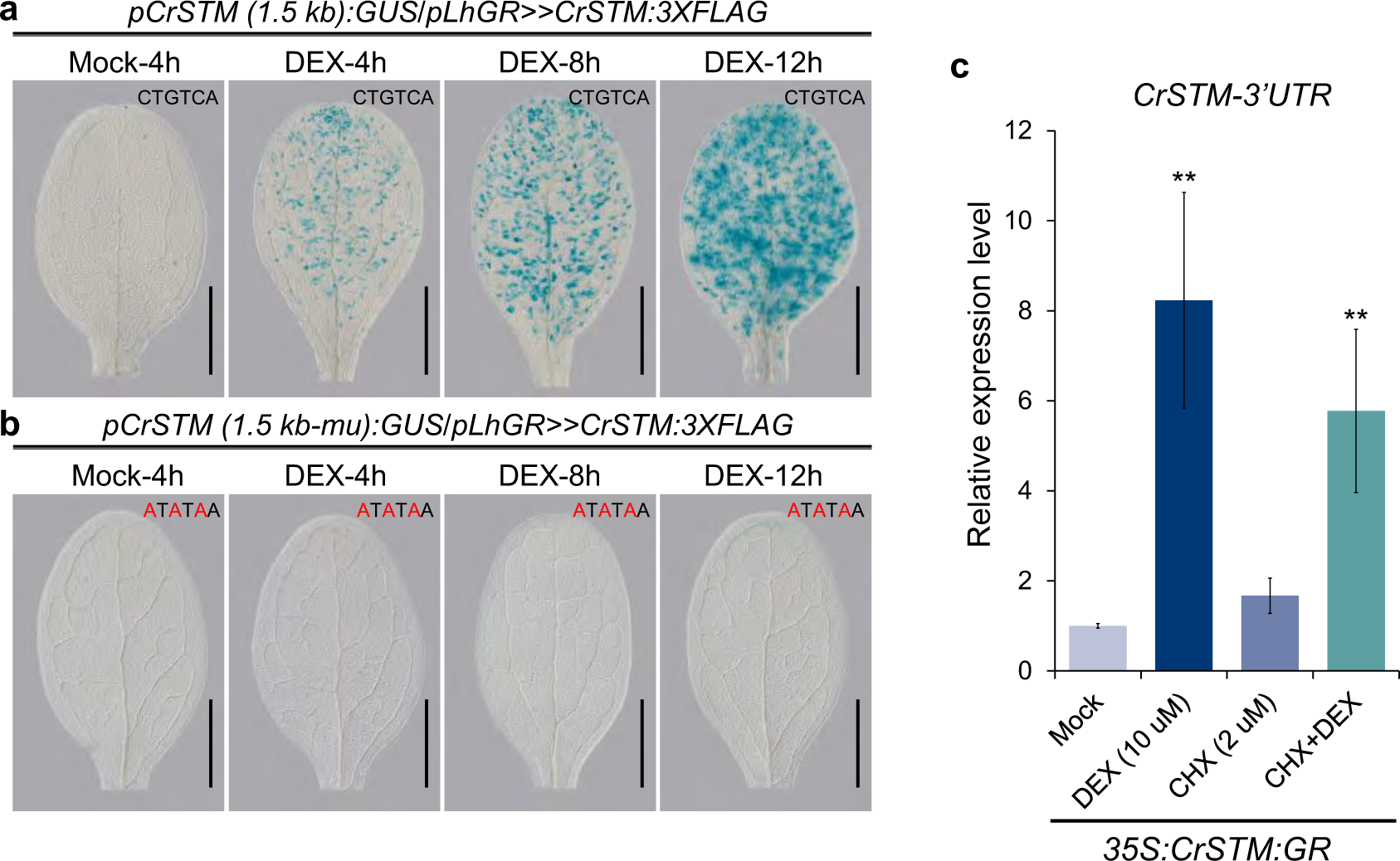
*CrSTM* directly activates its own expression. **a**, In the*pCrSTM (1.5kb):GUS/pLhGR>>CrSΊM:3xFLAG* lines, CrSTM:3*x*FLAG proteins is over-expressed upon DEX treatment but not in the mock at 4 hours’ time point. Over-expression of the CrSTM:3*x*FLAG proteins, in turn, activates the *pCrSΊM (1.5kb)* promoter shown by GUS staining in a time-course records. Please note that the expression of GUS is accumulated over time with DEX treatment, suggesting a response of *pCrSΊM (1.5kb)* promoter to the CrSTM:3*x*FLAG protein levels. **b**, In the *pCrSΊM (1.5kb-mu):GUS/pLhGR>>CrSΊM:3xFLAG* lines, *pCrSΊM (1.5kb-mu)* promoter does not respond to overexpression of CrSTM:3*x*FLAG proteins upon DEX treatment, suggesting the STM-binding site is responsible to the activation of GUS by CrSTM:3*x*FLAG. c, In the 35S:CrSTM:GR lines, endogenous *CrSΊM* expression (detected by probes targeting the 3’UTR) could be activated by CrSTM:GR proteins in the presence of DEX and protein synthesis inhibitor cycloheximide (CHX), indicating *CrSΊM* is a direct target of CrSTM:GR. Scale bars, **a** and **b**, 1 mm.

**Extended Data Fig. 17.**
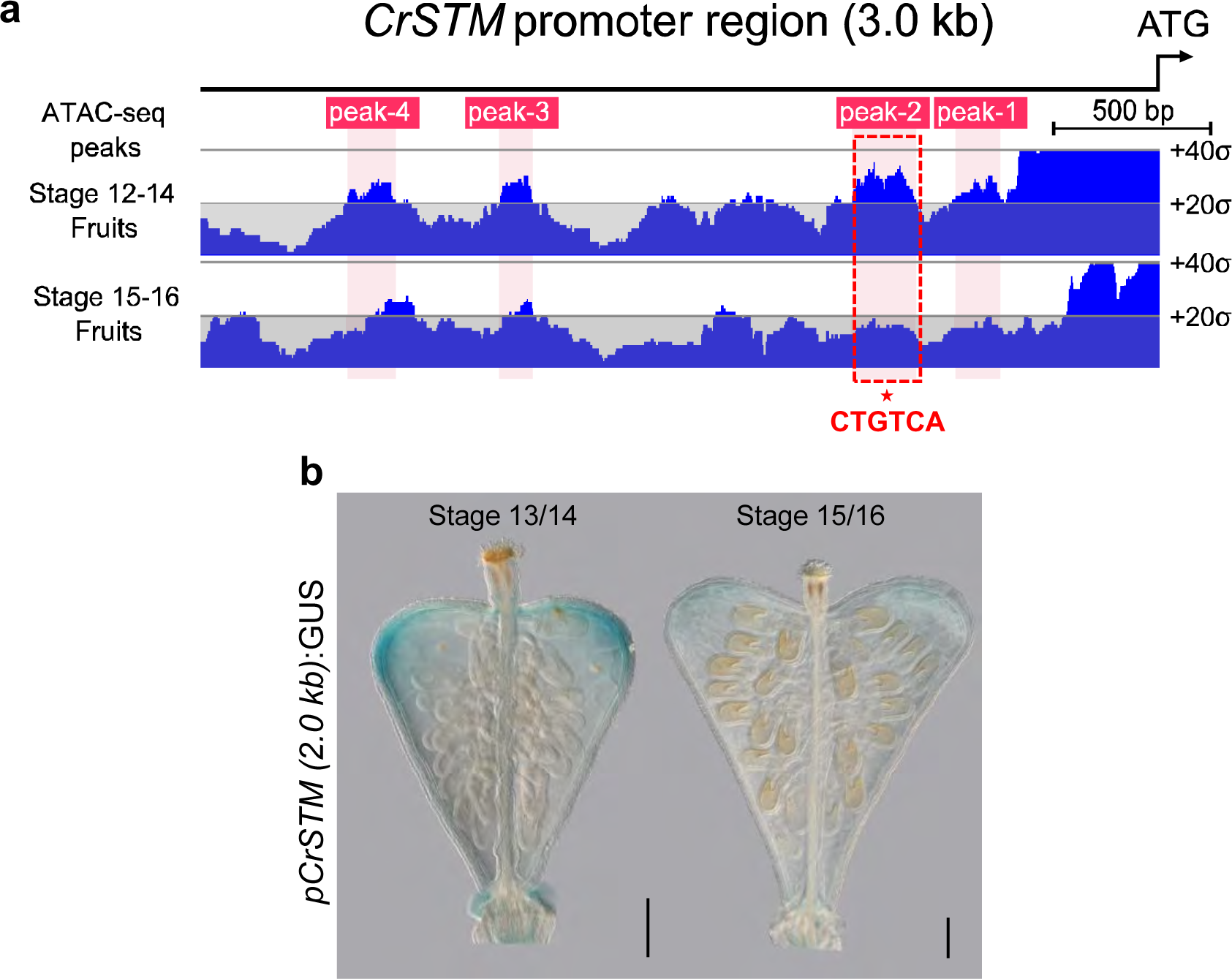
Expression of *CrSTM* is dynamically associated with the chromatin status in the STM-binding site. **a**, Assay for Transposase-Accessible Chromatin using Sequencing (ATAC-Seq) analysis of the *CrSΊM* promoter region (∼3.0 kb) in stage 12-14 fruits and stage 15-16 fruit samples. The grey lines mark the mean and standard deviation. ATAC-seq peaks are numbered sequentially from ATG and shaded in light pink. Please note that the chromatin region containing the STM-binding site, i. e. peak-2 (red-dash line boxed region), becomes less accessible in stage 15-16 fruit compared with stage 9-14 fruits, while other regions, such as peak-3 and peak-4 remain unchanged. **b**, Expression of CrSTM in stage 13/14 and stage 15/16 fruits shown by GUS staining in a *pCrSΊM (2.0 kb):GUS* reporter line. Scale bars in **b**, 100 μm.

**Extended Data Fig. 18.**
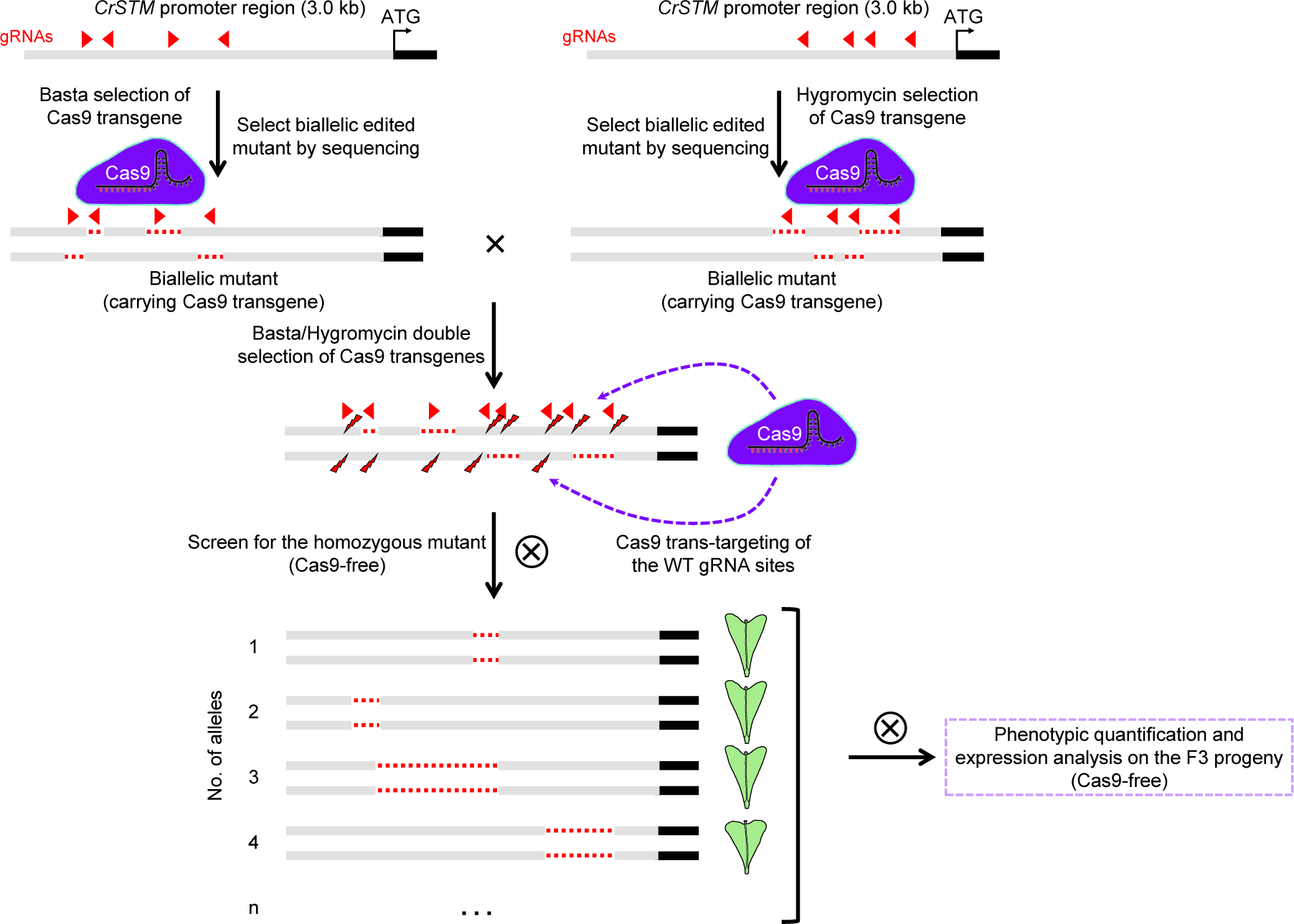
Schematic workflow generating the 22 *CrSTM* promoter mutant alleles using CRISPR/cas9. CRISPR/Cas9 transgenic plants are generated by transforming constructs carrying two sets of 4 gRNAs targeting the CrSTM promoter. Plants carrying the Cas9 transgene are screened by Basta and hygromycin (Hyg), respectively. The positive plants are then subjected to promoter genotyping, and those biallelic for promoter mutations are then crossed. The F_1_ plants inherited both Cas9 transgenes and associated 8 gRNAs (with both basta and Hyg selection marker) are screened and genotyped for biallelic mutations, confirming the effectiveness of Cas9 and gRNAs in gene editing. The F_1_ plants are selfed to generate a large segregation F_2_ population. The F_2_ plants are then screened simultaneously against Basta and Hyg (without Cas9 transgenes) on a plate and genotyped by PCR-sequencing for homozygosity for mutant alleles. The homozygous F_2_ plants of each genotype are selfed to the F_3_ generation for phenotypic and expression analysis. The promoter alleles in different F_3_ families are validated by PCR followed by sequencing. ATG indicates the start codon.

**Extended Data Fig. 19.**
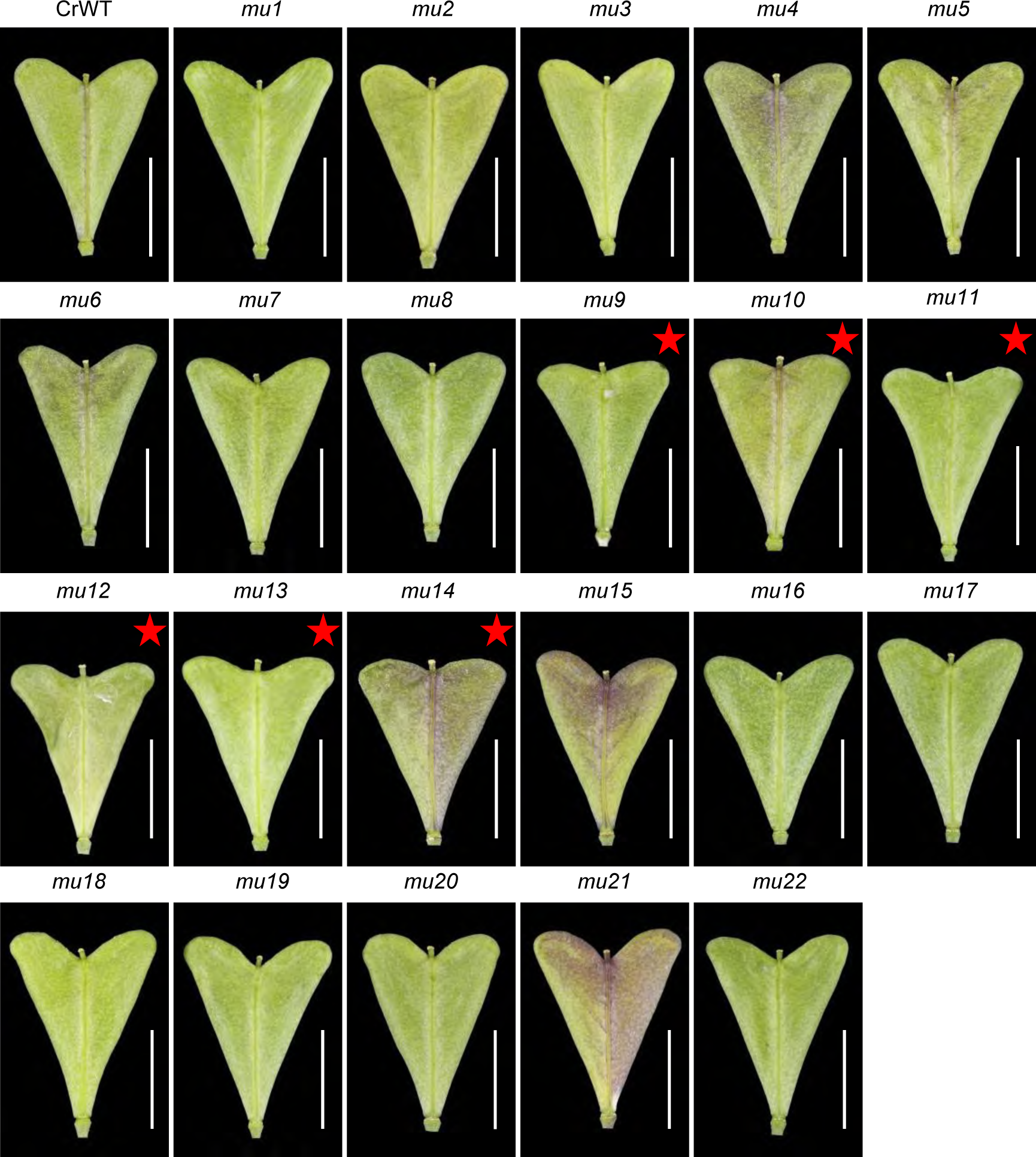
Fruit morphology of *CrSTM* promoter-edited alleles. Fruit morphology of the respective homozygous lines at stage 17. Each genotype corresponds to the details shown in Fig. 3k. Red stars indicate the mutant line with compromised out-growth of the fruit valve tips. Scale bars, 5 mm.

**Extended Data Fig. 20.**
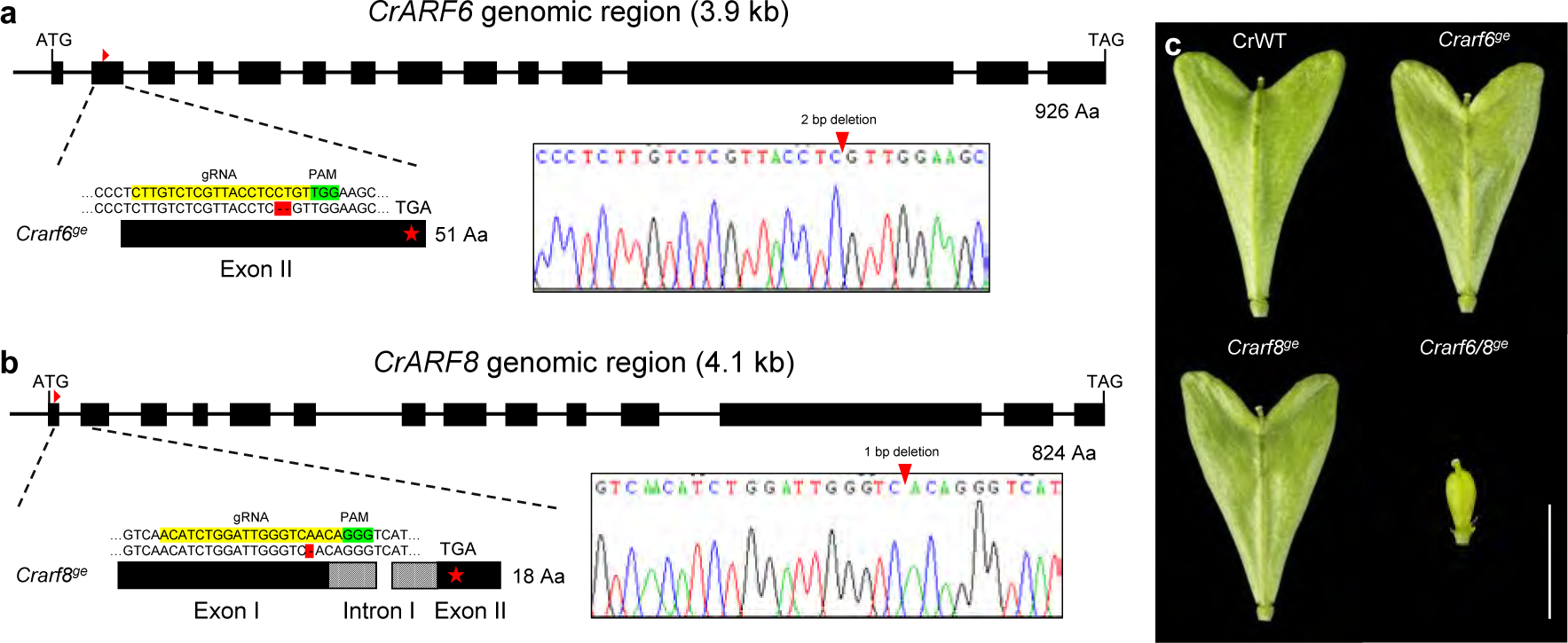
Generating the *Crarf6/8* mutant by CRISPR/Cas9. **a**, Schematic show of the *CrARF6* gene structure, the gRNA targeting the second exon is indicated by a red triangle. The *Crarf6^ge^* mutant is generated and confirmed by PCR-Sequencing of a 2-bp (CT) deletion in the second exon and the resultant protein is 51-aa in length compared with 926-aa in WT. **b**, Schematic show of the *CrARF8* gene structure. The gRNA targeting the first exon is indicated by a red triangle. The *Crarf8^ge^* mutant is generated and confirmed by PCR-Sequencing of a 1-bp (A) deletion in the first exon and the resultant protein is 18-aa in length compared with 824-aa in WT. **c**, Fruit morphology of WT, *Crarf6^ge^*, *Crarf8^ge^* and *Craf6/8^ge^* mutant at stage 17. The gRNA and PAM sequences are shaded in yellow and green, respectively. ATG and TAG indicate start and stop codons, respectively. The scale bar in **c** represents 5 mm.

**Extended Data Fig. 21.**
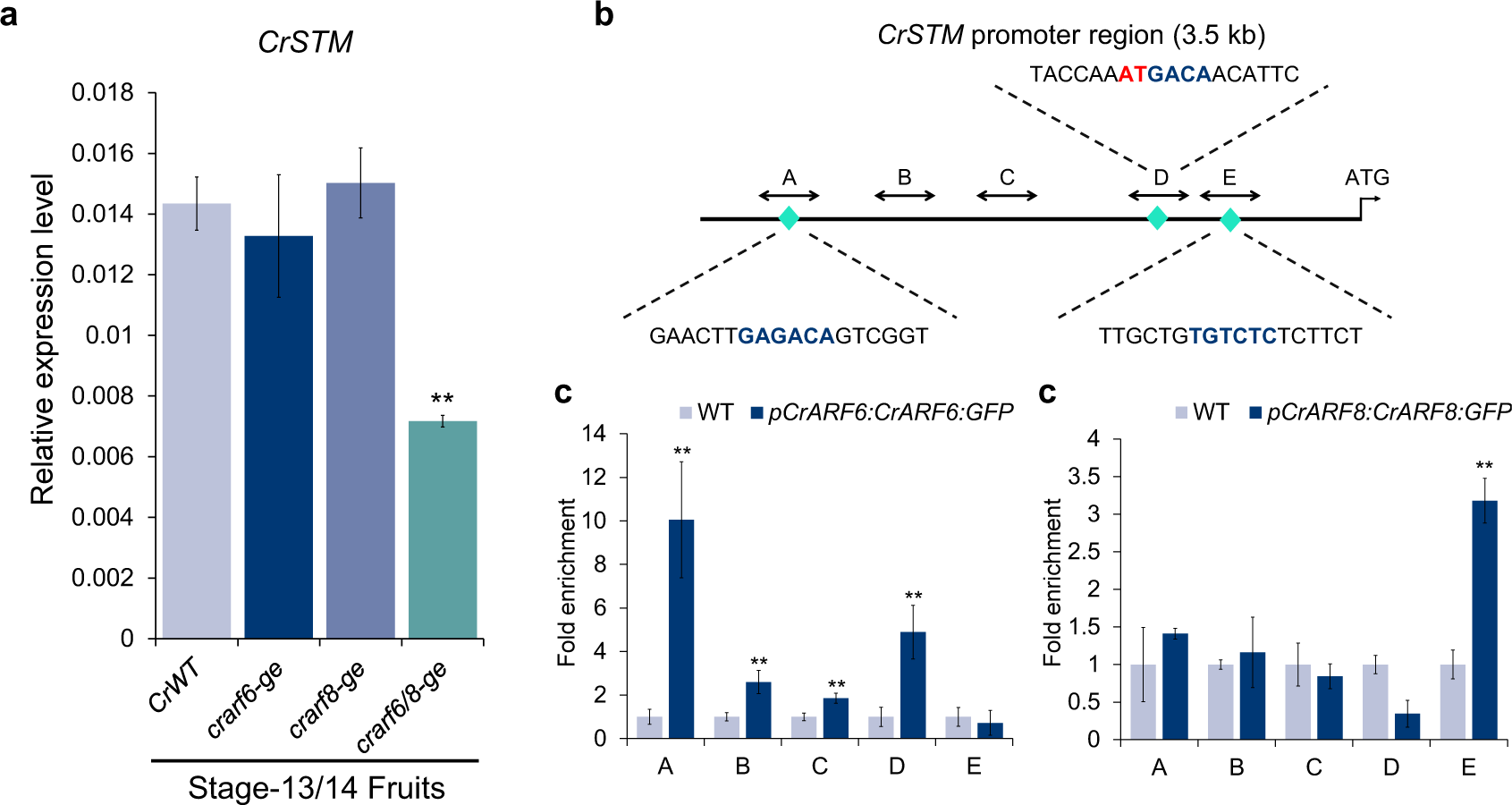
CrARF6/8 redundantly activate *CrSTM* expression in the fruits. **a**, In the stage 13/14 fruits, expression of *CrSTM* remain unchanged in the either *Crarf6^ge^* and *Crarf8^ge^* single mutant, while its expression is significantly decreased in the *Crarf6/8^ge^* double mutant, suggesting a redundant function of *CrARF6* and *CrARF8* in regulating *CrSTM*. **b**, Schematic show of the CrSTM promoter. The green diamonds indicate the auxin-responsive elements (blue letters). Double-head arrows indicate the regions subjected to ChIP-qPCR analysis. **c** and **d**, Chromatin Immuno­Precipitation (ChIP) analysis of CrARF6:GFP (**c**) and CrARF8:GFP (**d**) associated with the *CrSTM* promoter. Error bars represent the SD of three independent biological replicates. **, p < 0.01 (Student’s t-test).

**Extended Data Fig. 22.**
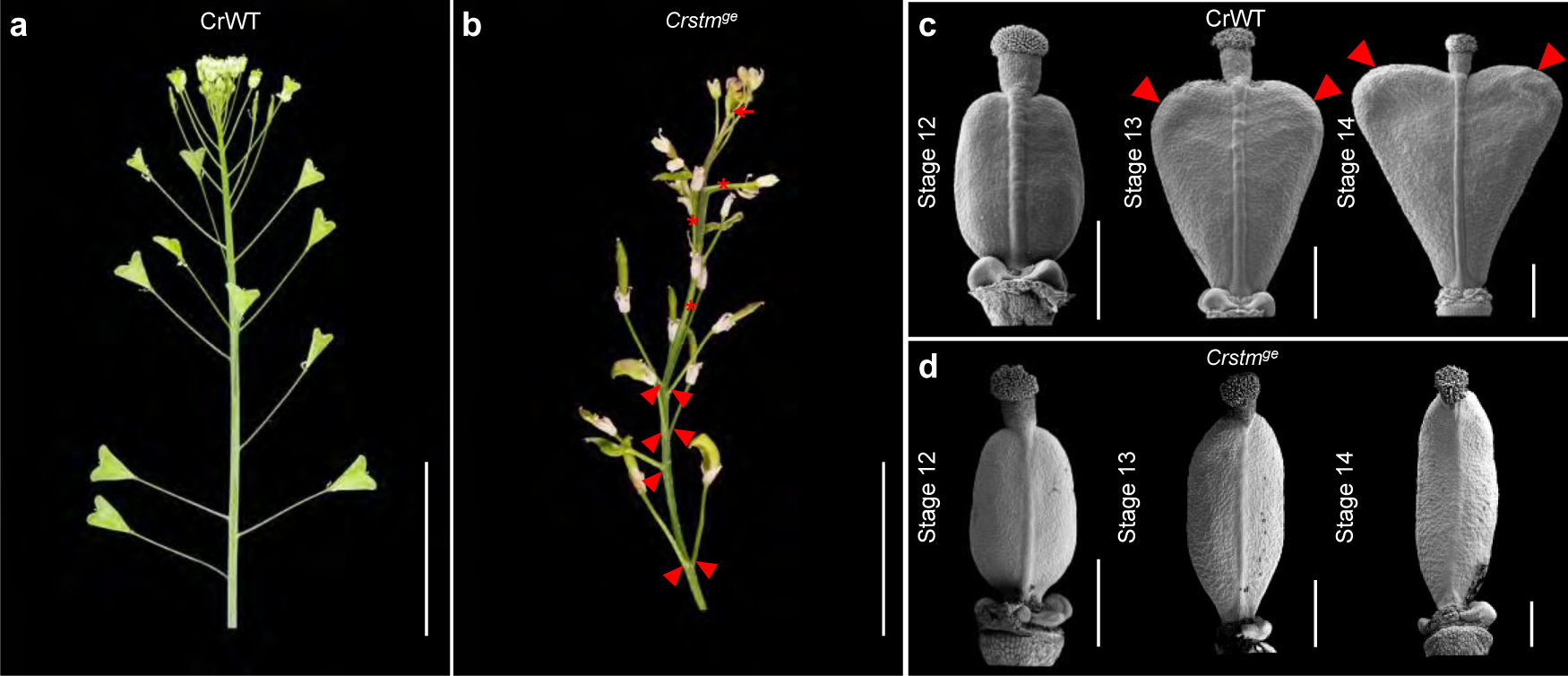
*Crstm* mutation disrupts the fruit shape transition process. **a** and **b**, Inflorescence phenotype of WT (**a**) and *Crstm^ge^* mutant (**b**). Please note that *Crstm^ge^* mutation generates pleiotropic defects in the inflorescence, including premature termination of the SAM (red arrow), fused pedicles (red asterisks) and altered phyllotaxis (red triangles). **c** and **d**, SEM analysis of WT (**c**) and *Crstm^ge^* (**d**) fruits from stages 12 to 14. Red triangles in **c** indicate the protrusions of the valve tips. Scale bars, **a** and **b**, 3 mm; **c** and **d**, 500 μm.

**Extended Data Fig. 23.**
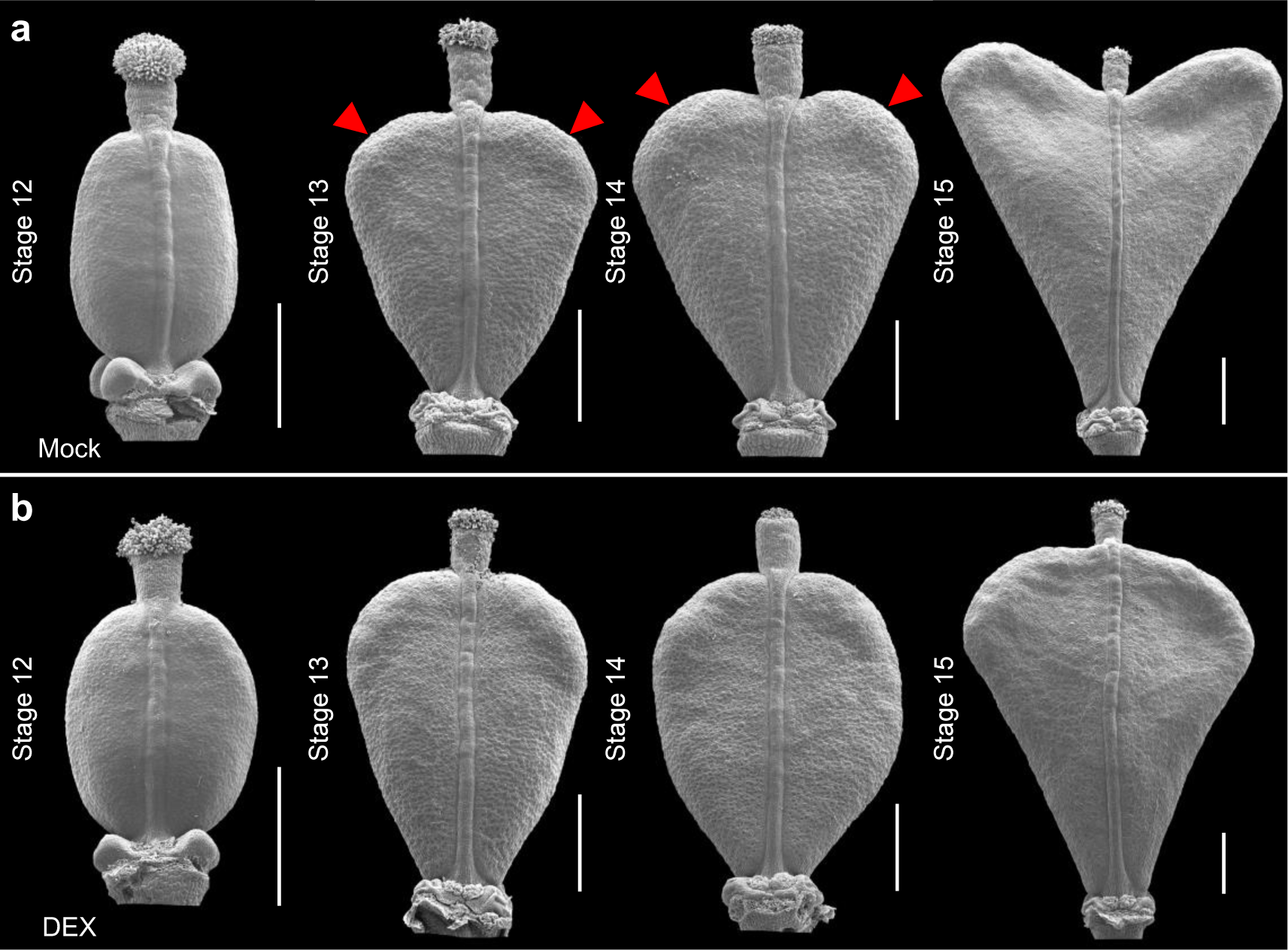
Down-regulation of *CrSTM* compromise the fruit valve tip development. **a** and **b**, SEM analysis of Mock (**a**) and DEX-treatment (**b**) fruits from stage 12 to 14 in the *pLhGR>>amiR-CrSTM* plants. Red triangles in **a** indicate the protrusions of the valve tips. Scale bars, **a** and **b**, 500 μm.

**Extended Data Fig. 24.**
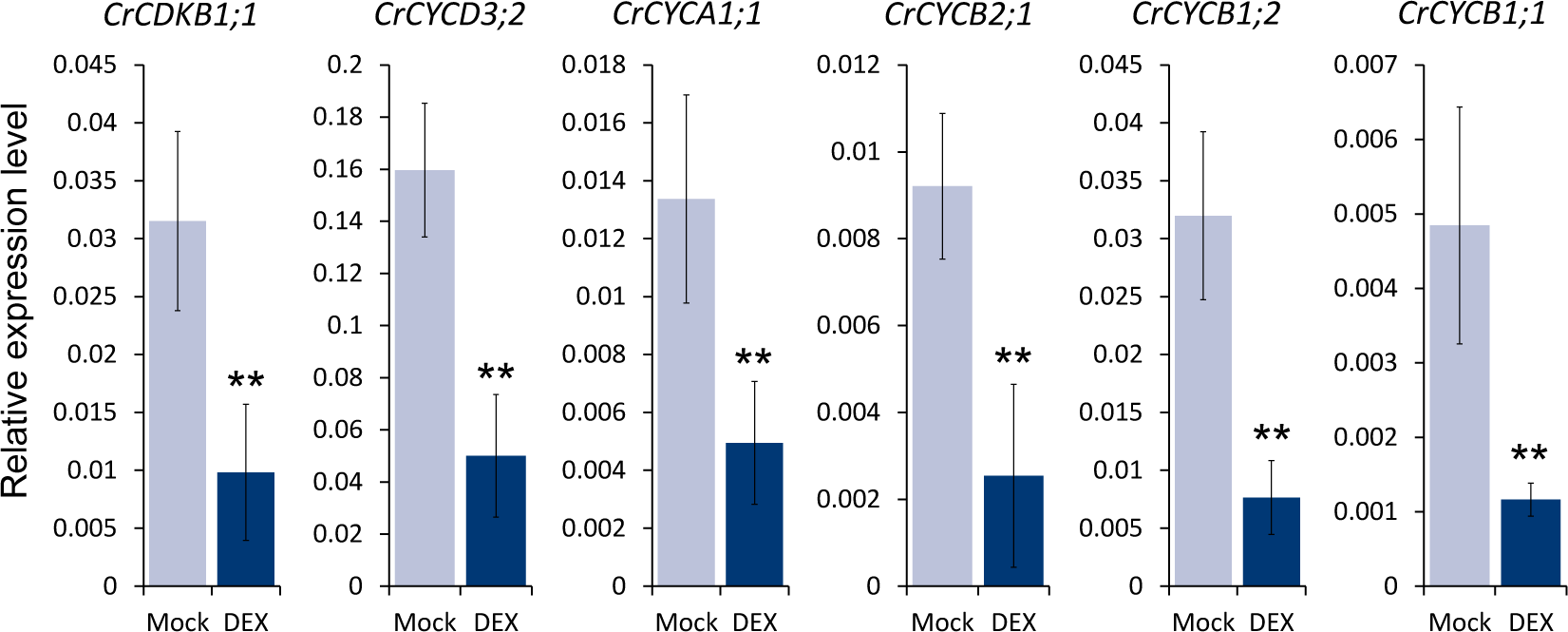
Expression analysis of cell cycle genes in the *pLhGR>>amiR-CrSTM* lines. qRT-PCR analysis of genes involved in the cell cycle shows that these genes are significantly down-regulated upon DEX treatment. Error bars represent the SD of three biological replicates. **, p < 0.01 (Student’s t-test).

**Extended Data Fig. 25.**
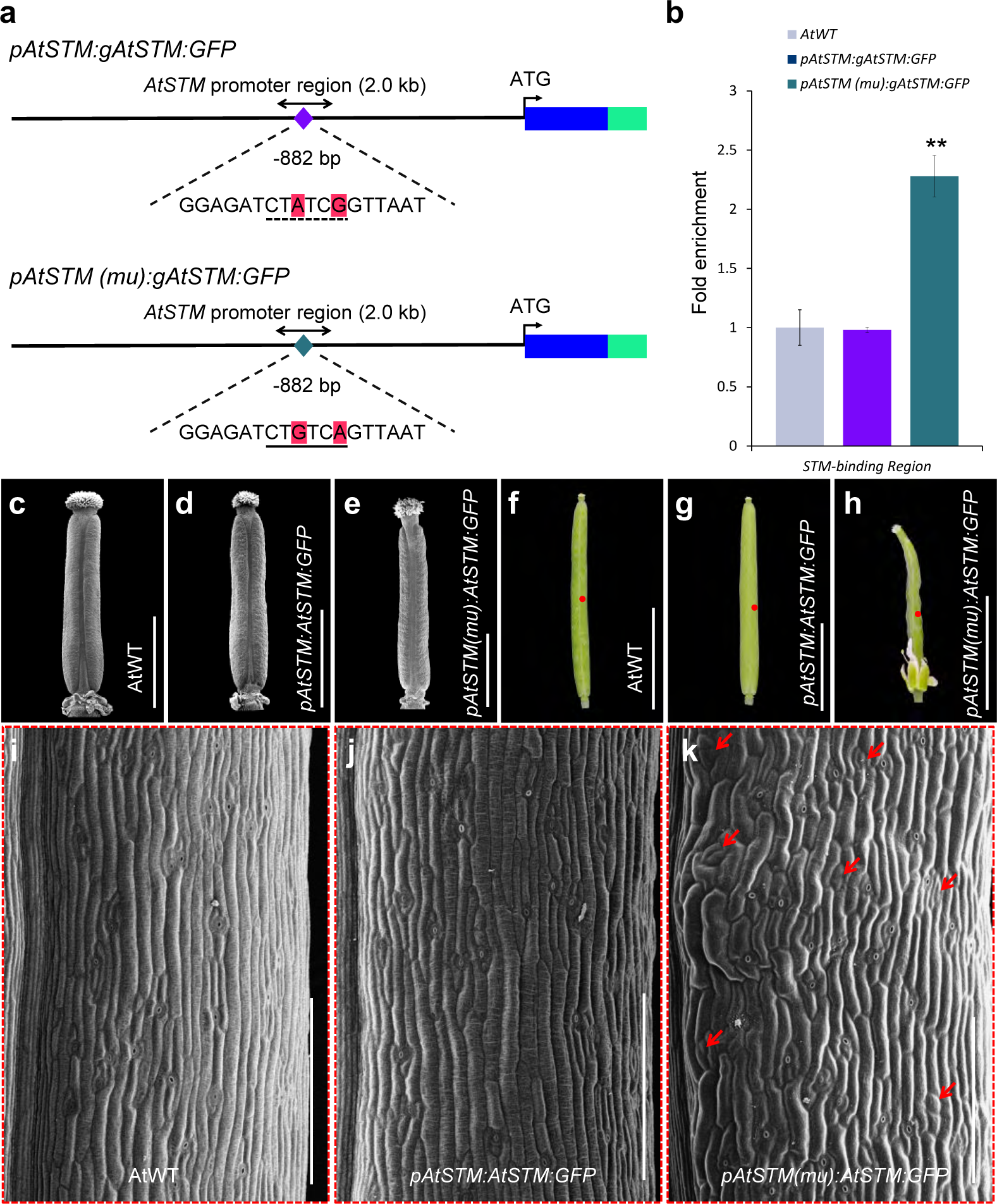
Gain-of-function mutation of the STM-binding site in *AtSTM* promoter disrupts fruit development. **a,** Schematic show of the *in situ* gain-of-function mutation of the STM-binding site in the AtSTM promoters. The different base-pairs were shaded in red; the STM-binding site was underlined with a solid line and the WT counterpart was underlined with a dash line, respectively. Please note that in the *pAtSTM (mu):AtSTM:GFP* construct, the native “CTATCG” sequences are mutated into “CTGTCA” STM-binding site, while the other promoter sequences remain unchanged. The length of each gene is not to scale. **b**, Chromatin Immuno-Precipitation (ChIP) analysis of AtSTM:GFP associated with the *AtSTM* promoter (shown in double arrows in **a**). Error bars represent the SD of three independent biological replicates. **, p < 0.01 (Student’s t-test).**c**-**e**, SEM images of the stage 11 gynoecia from AtWT (**c**), *pAtSTM:AtSTM:GFP* (**d**) and *pAtSTM (mu):AtSTM:GFP* (**e**) plants. **f**-**h**, Fruit morphology at stage 17 of AtWT (**f**), *pAtSTM:AtSTM:GFP* (**g**) and *pAtSTM (mu):AtSTM:GFP* (**h**) plants. **i**-**k**, SEM images of the epidermal cells from the fruits of AtWT (**i**), *pAtSTM:AtSTM:GFP* (**j**) and *pAtSTM(mu):AtSTM:GFP* (**k**), corresponding to the red-dotted region shown in the respective genotype in **f** to **h**. The arrows in **k** indicate the cells are still ongoing cell division. Scale bars, **c**-**e**, 1 mm; **f**-**h**, 5 mm; **i**-**k**, 200 μm.

**Extended Data Fig. 26.**
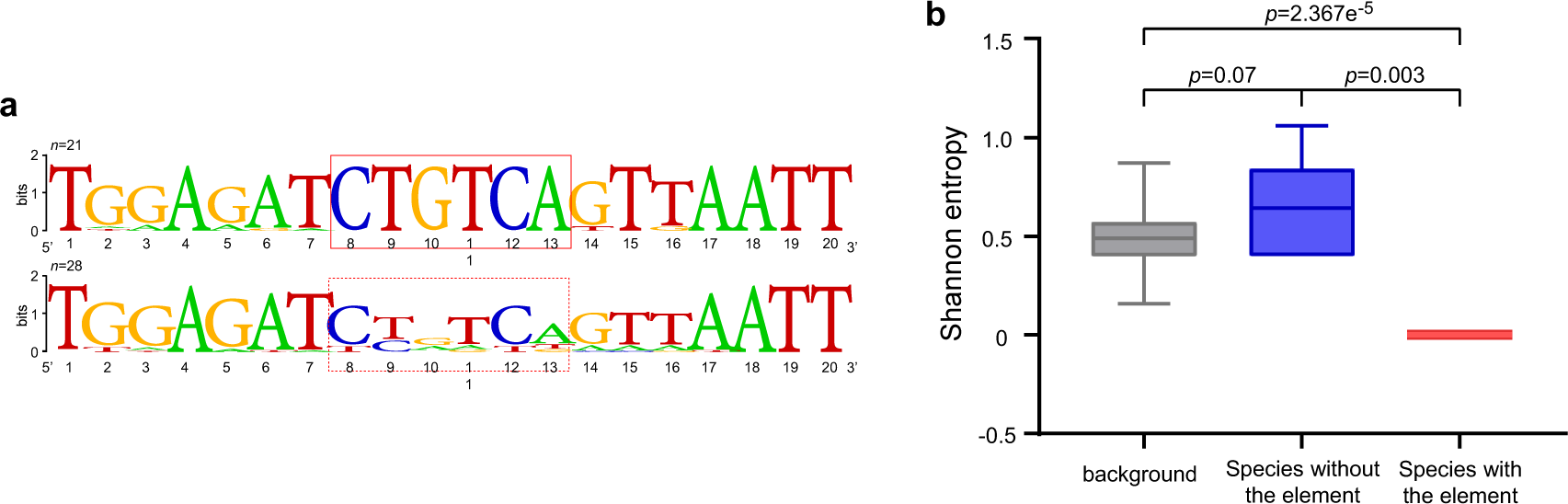
Detection of strong selective constraint on the STM-binding site in Brassicaceae. **a**, Comparison between the sequence logos of the STM-binding site from species evolved with this element (upper panel, n=21) and without this element (lower panel, n=28). Please note that the STM-binding site is 100% identical among the species that evolved with this element, while it is less conserved in the counterpart region among species without this element. The plot is generated using WebLogo online tool (https://weblogo.berkeley.edu/). **b**, Conservation comparison between the STM-binding site and its counterpart, using selectively neutral sites as control. Shannon entropy of the 6-bp element and its counterpart for each species was calculated and grouped accordingly. Neutral sites are four-fold degenerate sites (4-fold sites) from STM-orthologous genes. We generated the control group by randomly sampling six sites from all the 4-fold sites each time with 1000 repeats. *p*-values are from the two-sided Mann-Whitney U test.

